# Protein Language Model Supervised Scalable Approach for Diverse and Designable Protein Motif-Scaffolding with GPDL

**DOI:** 10.1101/2023.10.26.564121

**Authors:** Bo Zhang, Kexin Liu, Zhuoqi Zheng, Junjie Zhu, Zhengxin Li, Yunfeiyang Liu, Junxi Mu, Ting Wei, Hai-Feng Chen

## Abstract

Proteins perform essential roles in numerous biological processes, largely driven by the three-dimensional structure of several key motif residues. Recently, a variety of energy-based and machine learning backbone generation methods have been developed to solve the motif-scaffolding task. However, it is still challenging to generate diverse and accurate scaffold structures around motifs for models either fine-tuned pre-trained multiple sequence alignment-based (MSA-based) structure prediction models or trained from scratch. Here, we introduced Generative Protein Design by Language model (GPDL) for effectively replacing traditional MSA-based pretraining. Using our scalable design strategy, GPDL successfully solved 22 out of 24 benchmark problems and outperformed other methods by generating 33.5% more unique designable clusters than RFdiffusion. This demonstrates that our approach can generate accurate and physically plausible structures across diverse protein design scenarios. GPDL also showed strong robustness in orphan proteins that have low sequence similarity with the training set. Our approach underscores the promise of protein language models in protein design and has the potential to accelerate the discovery of novel functional proteins for a wide range of biological and therapeutic applications.

## Introduction

Protein design has advanced significantly in recent years, especially with deep learning breakthroughs like AlphaFold2^1^. Many of these approaches follow a motif-scaffolding paradigm, which enables the creation of functional novel proteins, including enzymes^2^, binders^3^, and antibodies ^4^. Within this approach, functional motifs are defined as specific spatial arrangements of amino acid segments, and these works generate new protein backbones to accurately position these regions in their original place.

The primary challenge in motif-scaffolding lies in generating diverse yet consistent scaffolds while preserving the original motif 3D structures with sub-angstrom accuracy. Achieving this requires stabilizing interactions both within and outside the motif regions. Current state-of-the-art methods typically address this challenge by fine-tuning pretrained protein structure models, such as RFdiffusion ^5^ and RFjoint ^6^. While these approaches have significantly improved design accuracy, they often lead to low diversity in scaffolds. This limitation may arise from the inability of MSA-based pretrained models to effectively sample the structural space when fine-tuned for de novo designed proteins, which typically lack co-evolutionary information. Other methods trained from scratch are capable of generating diverse structures to some extent, while often at the cost of lower accuracy^7,8^. On the other hand, several approaches have also explored the use of Markov Chain Monte Carlo (MCMC) or gradient-based algorithms for *de novo* protein design, typically referred to as “hallucination-based” approach, such as restrained hallucination ^6^, relaxed sequence optimization ^9^ and BindCraft^10^. However, none of these approaches have yet applied language models as prompts for generating functional proteins.

To address this challenge, we leveraged protein language models, trained solely on amino acid sequences without homologous information, to effectively sample the protein space. Protein language models are large pre-trained models utilized for sequence representation^11–14^ which can be fine-tuned for various downstream tasks, including protein structure prediction ^15,16^, *de novo* sequence generation^17–19^, protein function annotation, and protein-protein interactions ^20^. These models inherently capture long-range dependencies within protein sequences due to their extensive parameters and large-scale training datasets.

We initialized a structure seeding module with a fine-tuned structure module adapted from ESMFold ^15^, resulting in a prompt structure. ESM-IF1 ^21^ was later applied to encode this structure into possible sequences. These sequences were then fed into a structure optimization module, refining sequences through a MCMC-based simulated annealing schedule. Our seeding and optimization module offers enhanced efficiency compared to optimization methods based on purely random sequences. The results demonstrated that our model solved more problems and generated more unique designable clusters compared to other methods.

## Results

### Overview

In this study, we used the protein language model ESM2 along with the folding trunk and structure module from ESMFold to develop GPDL (Generative Protein Design by Language model). GPDL consists of two main components: the structure seeding module, which identifies an optimal prompt structure; and the structure optimization module, which iteratively refines this into a highly plausible structure. The model architecture is illustrated in **Fig. 1**, with training details provided in the materials and methods.

**Figure 1.**
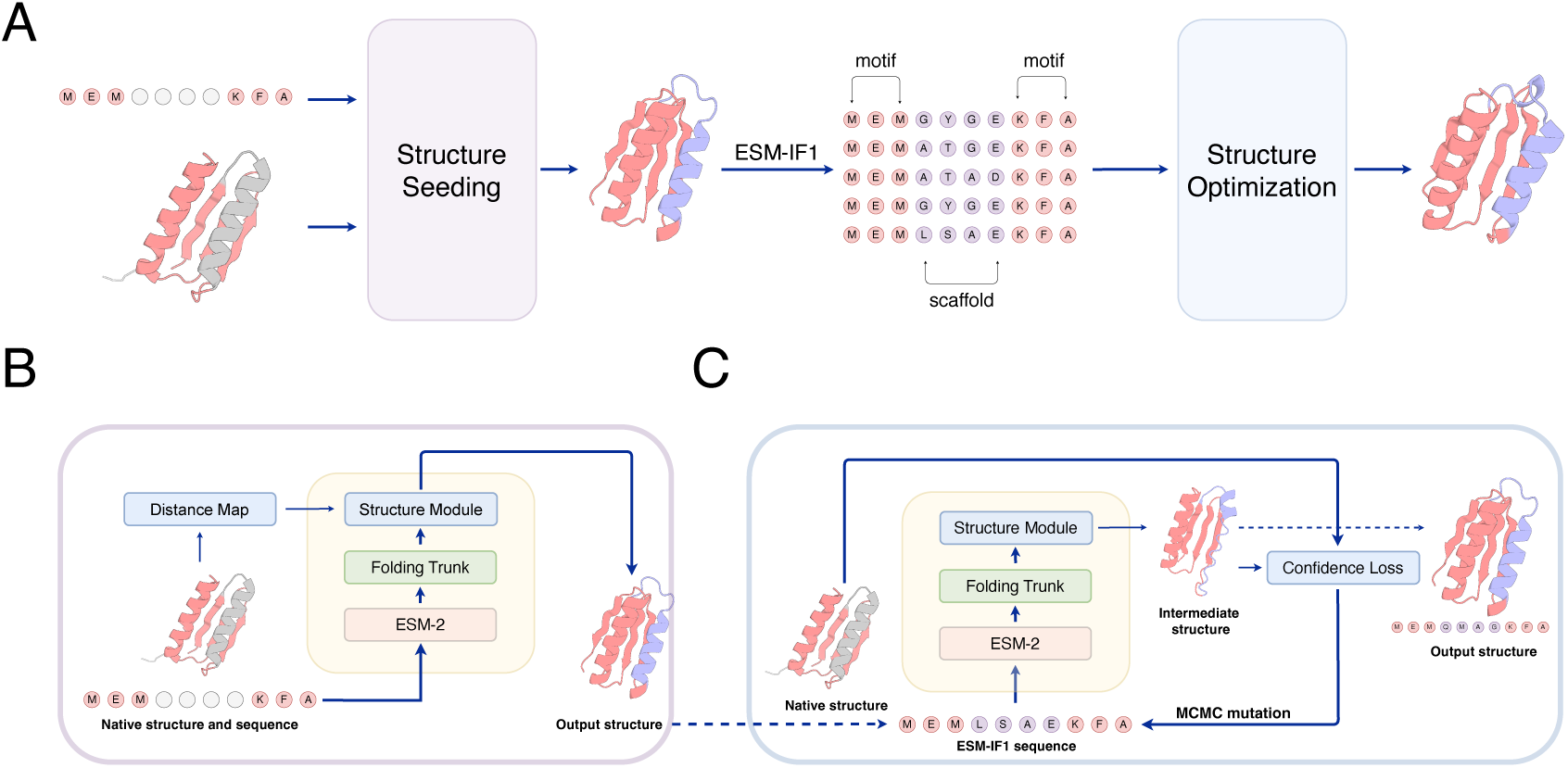
Model architecture. (A) The GPDL workflow. (B) The structure seeding model. (C) The structure optimization model. The input structure on the left has the motif region (pink) and scaffold region (gray). The output on the right is a designed structure with the conserved motif (pink) and a newly designed scaffold (blue). The PDB ID in the plot is 2KL8.

The structure seeding module (**Fig. 1B**) takes as input a specified motif sequence and its distance map to produce an initial, prompt structure, which is subsequently encoded into sequences using ESM-IF1. This module was fine-tuned under an inpainting task, and we extended the original architecture, enabling the retrieval of motif information and facilitating the generation of corresponding structures and sequences. The structure optimization module (**Fig. 1C**) was then utilized to iteratively refine this sequence under the supervision of loss functions regards to structure deviation and prediction confidence, which finally resulted in a highly optimal scaffold.

### Accurate and scalable motif-scaffolding with GPDL

To rigorously evaluate GPDL’s performance, we used a benchmark set covering diverse motif-scaffolding cases from RFdiffusion ^5^. We additionally compared the performances along with other outstanding methods including RFdiffusion ^5^, RFdesign ^6^, Chroma ^8^, and TDS ^22^. GPDL successfully solved 22 out of the 24 cases (**Fig. 2A**), achieving the highest success rate among all tested methods. Notably, GPDL reached at least the same level of accuracy compared to RFdiffusion but was capable of generating more diverse structure clusters (**Tables S1-S2**).

**Figure 2.**
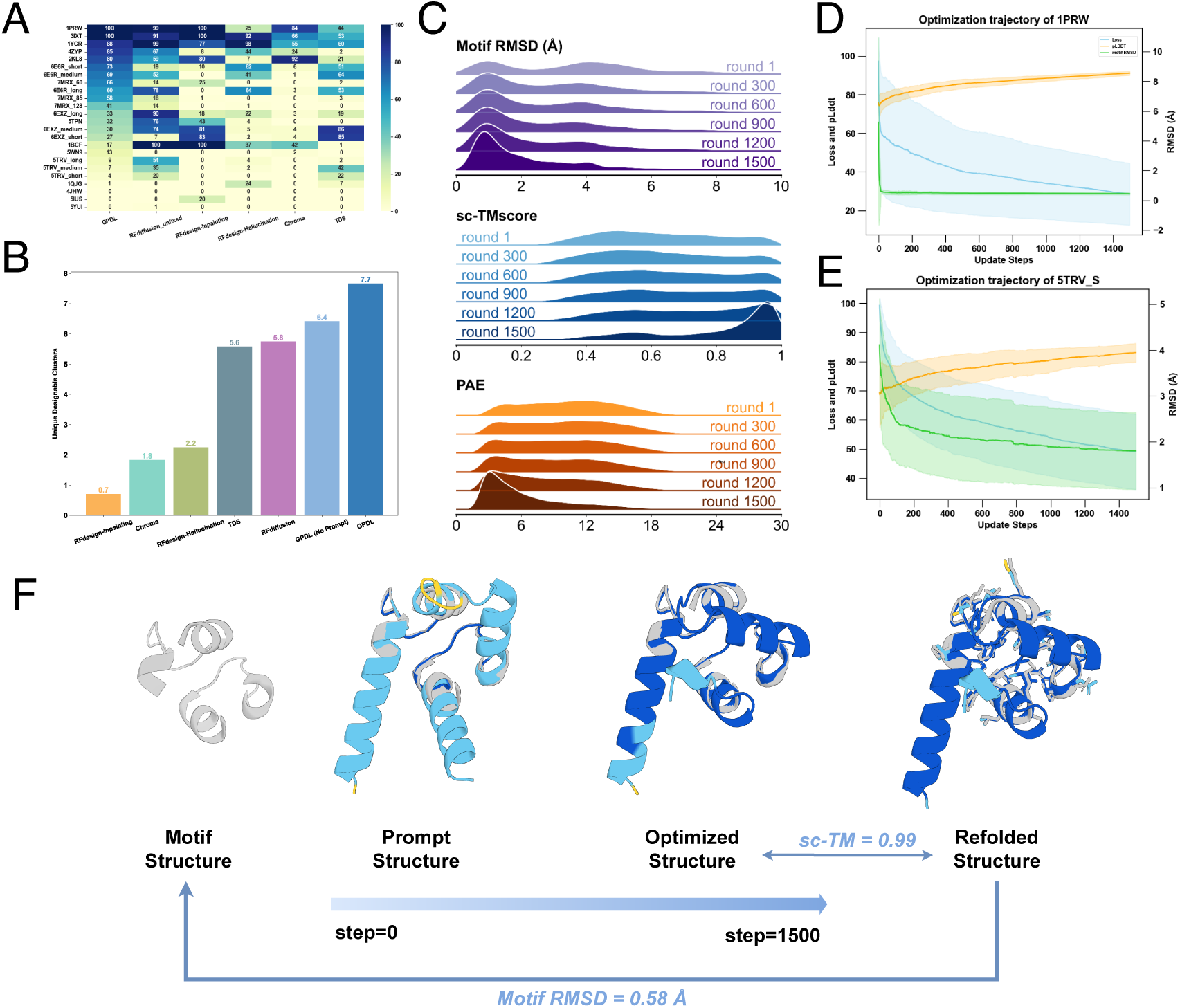
Overall performance. (**A**) Success rates of 24 benchmark cases from each method. (**B**) Unique designable clusters for each method. The attached numbers denote the number of clusters within designable backbones averaged across the whole benchmark. (**C**) Optimization trajectories in different steps, with metrics representing the mean values of the backbones during optimization: top, motif RMSD; middle, sc-TMscore; bottom, PAE. (**D-E**) Loss curves for benchmark cases 1PRW and 5TRV (short) during optimization. (**F**) Design pipeline from an initial motif structure to a prompt structure generated by seeding module, followed by an optimized structure with 1500 MCMC mutations. The displayed structure is based on the 1PRW motif. The final refolded structure has a motif RMSD of 0.58 Å relative to the native motif, an sc-TMscore of 0.99 compared to the designed structure, pLDDT of 90.66 and max PAE of 2.43. The designed and refolded structure are coloured by the pLDDT confidence score, with blue in high-confidence regions.

In order to consider the designability and diversity in the meantime, we used unique designable clusters ^23^ to quantify the variety of successful design clusters, which better reflects the generalization of the models and the extension of protein space beyond measuring designability and diversity separately. Under this scenario, GPDL generated the highest number of unique designable clusters among all methods (**Fig. 2B**), with a 33.5% increase over RFdiffusion, underscoring its capacity to produce a diverse range of viable designs. Results further showed that the seeding module can provide prompt structures with greater diversity (**Table S1**). Notably, when no prompt structures were introduced, GPDL generated less unique designable clusters, underlining the significant role of the seeding module. We hypothesize that this might arise from the inherent features of protein language models trained using an enormous number of protein sequences, while the MSA-based models trained with limited protein crystal structures potentially lack this prior knowledge of protein space.

While GPDL benefits from the seeding module in terms of diversity, the structure optimization module plays a crucial role in generating more refined and confident backbones, improving both success rates of design and computational efficiency with greater scalability of GPDL. Key metrics, such as motif RMSD, self-consistency TMscore (sc-TMscore), and PAE, demonstrated consistent convergence throughout our optimization trajectories. We noted that metric converged at a significantly different speed with respect of the motif hardness. For challenging cases like a curved beta-sheet protein (PDB ID: 5TRV), as optimization rounds increased, we observed a shift in a subset of the prompt protein designs given by the seeding module towards more favorable outcomes, evidenced by a peak in higher TMscore and lower PAE values, and a gradually convergence in several key metrics (**Fig. S1 and Fig. 2C**). This indicates that the optimization module effectively refined the initial prompt structures toward optimal results (**Fig. S2 and Fig. S3**).

In the previous methods, scaffolding the active site KSI, a catalytic triad composed of an aspartic acid, an asparagine, and a tyrosine (PDB ID: 1QJG) presents also challenging due to its conformational flexibility and extremely limited number of motif residues (**Table S2**). Optimization module is crucial for accurately reconstructing this catalytic triad with sub-angstrom accuracy, maintaining the orientation of the side-chains where other methods such as RFdiffusion and RFdesign-inpainting failed to scaffold it. It’s worth mentioning that the generated backbone adopts a fold distinct from the native reference, positioning aspartic acid and tyrosine in α-helix, whereas they reside in α-helix and β-sheet in the native structure. The loop supporting asparagine remains intact, which, given the loop’s high flexibility, helps to retain the catalytic potential of the designed protein (**Fig. S4**). Moreover, the reduced backbone size compared to the reference highlights the potential of our method to search for smaller functional protein backbones, a promising feature for applications in medical research and healthcare.

In simpler cases, such as the EF-hand motif with a helix-loop-helix secondary structure (PDB ID: 1PRW), fewer optimization rounds were sufficient to achieve favorable results, accelerating the design process and reducing computational costs while maintaining high-precision, reliable protein structures (**Fig. 2D and Fig. S1**). In contrast, harder cases required more iterations to reach convergence (**Fig. 2E**). For previously challenging cases where GPDL default had failed—such as the PD-L1 with a binding interface on PD-1 (PDB ID: 5IUS), characterized by a two-segment, double-layer beta-strand motif—RFdiffusion generated only one successful design, while our method produced four successful designs with an iteration count of 5000 (**Fig. S5**).

The user-specified iteration cycles enhance both the scalability and efficiency of the method. In ablation experiments, the seeding and optimization modules independently performed well, accurately reconstructing the motif and overall backbones (**Fig. S4**). Integrating these modules enables GPDL to achieve an optimal balance between design accuracy and computational efficiency (**Fig. 2F**).

### Designed Backbones of GPDL showed diverse and reasonable topologies

We further analyzed the designs in each case, focusing particularly on diversity, stability, and other metrics relevant to designability. We randomly sampled 50 successful backbones from both GPDL and RFdiffusion across a same benchmark case with different lengths. Diversity was quantified through multi-dimensional scaling (MDS) based on pairwise backbone TMscore among successful designs. The pairwise TMscore distribution for GPDL shifted to the left with a lower peak, indicating greater diversity in successful designs (**Fig. 3A and Fig. 3D**). And the MDS plot for GPDL covered a broader range of topologies compared to RFdiffusion (**Fig. 3B and Fig. 3E**). These results suggest that, although RFdiffusion has a higher success rate, GPDL is more likely to explore a wider structure space.

**Figure 3.**
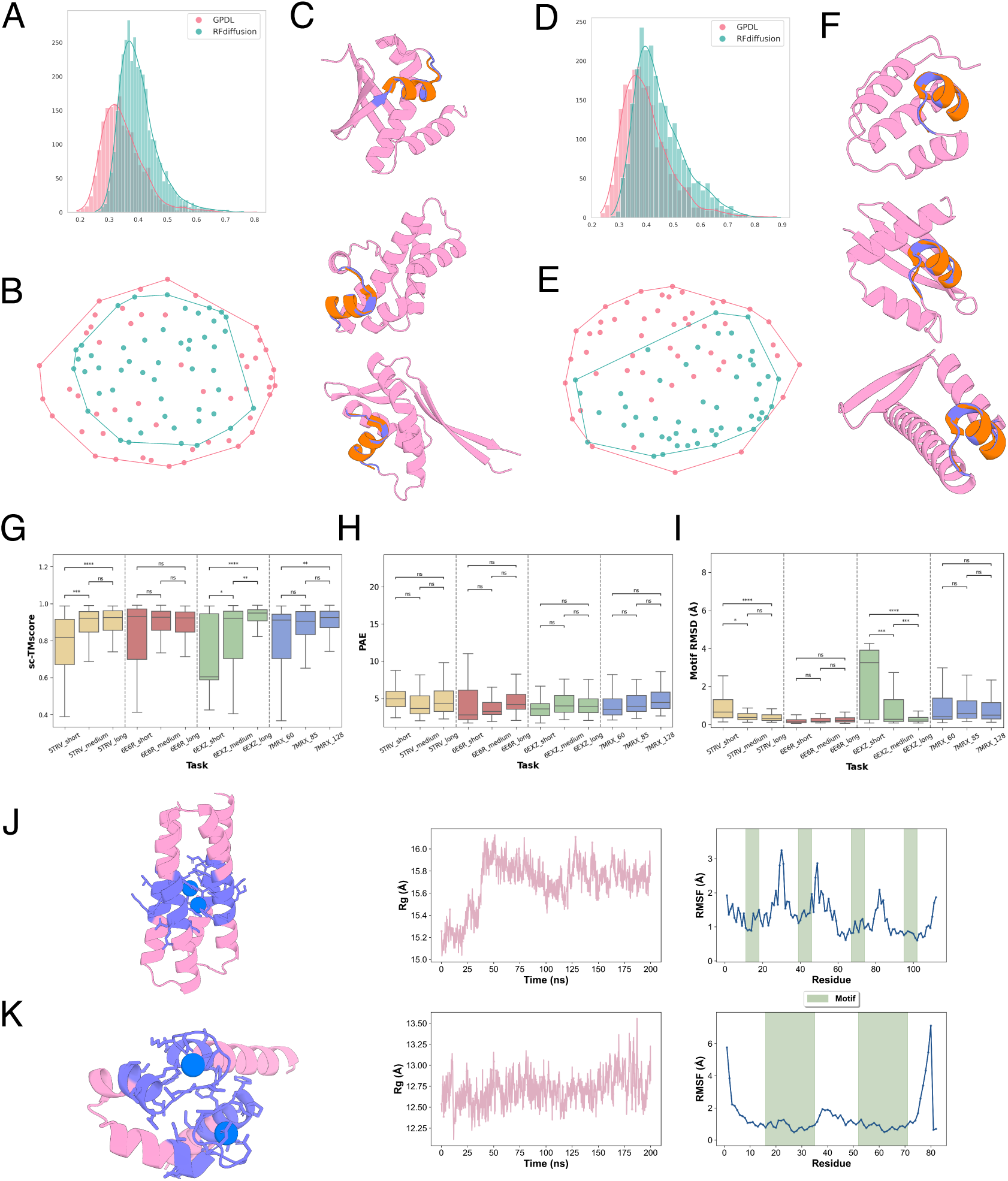
GPDL generates diverse and physically reasonable structures for different lengths. (A-C) Results for 6E6R_long: (A) Pairwise TMscore distribution, (B) MDS plot of pairwise TMscore, (C) Visualizations of designed backbones exhibiting distinct topologies. (D-F) Results for 6E6R_medium: (D) Pairwise TM-score distribution, (E) MDS plot of pairwise TMscore, (F) Visualizations of designed backbones with varied topologies. (G-I) Comparative metrics across lengths: (G) sc-TMscore, (H) PAE score, and (I) motif RMSD. (J-K) AlphaFold3 predictions and MD simulations for selected designs. The AF3-predicted structures are shown on the left, where fluctuations during simulation are displayed on the right. The statistics are collected from MD simulation with 200 ns. (J) Results for 1BCF. (K) Results for 1PRW. Colors of protein structures: Native functional motifs aligned to designed structure, orange; designed scaffold, pink; designed motif, purple; predicted ions, blue.

The ability to design optimal short scaffolds is especially valuable as it suggests potential for creating functional proteins with smaller backbones, which is significant for applications in drug delivery and therapy (**Fig. 3G-I**). GPDL demonstrated robust performance in designing scaffolds across various lengths, particularly for the barnase ribonuclease inhibitor protein (PDB ID: 7MRX), where it consistently outperformed RFdiffusion regardless of scaffold length. For the ferredoxin protein (PDB ID: 6E6R) long and medium scaffold tasks, both GPDL and RFdiffusion performed comparably, but in short scaffold tasks, GPDL held a notable advantage. For the RNA export factor (PDB ID: 6EXZ) motif, GPDL’s performance varied with scaffold length, likely due to the helix-loop-sheet topology of motif, which may require longer scaffolds for adequate support (**Fig. S6-S11**). With extended scaffolds, GPDL also achieved higher design diversity, with the resulting structures covering a broader range of optimal backbones.

Metal-binding proteins exhibit a broad distribution in biological systems and perform significant roles across numerous biological processes ^24^. Thereupon, we investigated the design of di-iron binding protein (PDB ID: 1BCF) and a double EF-hand motif with two calcium ions (PDB ID: 1PRW). We first utilized ProteinMPNN ^25^ to generate sequences compatible with backbones designed by GPDL and used AlphaFold3 (AF3)^26^ to predict three-dimensional structures with corresponding ions. The resulting structures showcased that the designed motifs successfully hold metal ions with stable binding interactions (**Fig. S12**). In order to further explore the stability of designed motifs, we performed all-atom molecular dynamics (MD) simulations upon these structures and analyzed the fluctuations of key biophysical properties during simulations. Rg (Radius of Gyration) reflects the compactness of simulated structures and is a commonly-used metric to indicate the overall stability of protein structures during MD simulation. Our results show a tendency of convergence after 200 ns MD simulation, demonstrating the stability of the overall structures. Additionally, the small fluctuations of motif regions could be observed from the results of residue-wise RMSF (Root Mean Square Fluctutation), further suggesting the stability of metal-binding motifs, which exhibit significant relevance to biological applications of metal-binding proteins.

### Model showed strong robustness in low homologous sequences

To test the generalizability of GPDL, we used an orphan protein benchmark set derived from RFdiffusion ^5^. Among these 30 proteins, we classified them into two groups: the orphan group, with a maximum sequence similarity of less than 30% to the training sequences using MMseqs2 ^27^, and the training group, which includes the remaining proteins. Results obtained using the same evaluation pipeline indicate robust performance across both orphan and training proteins, with comparable outcomes in global TMscore, PAE, motif RMSD, and pLDDT (**Fig. 4A-D)**. Our model demonstrated strong robustness to proteins with low sequence similarity to those in the training set. Notably, orphan proteins achieved even higher success rates than training proteins (**Fig. 4E**), aligning with findings from RFdiffusion’s orphan protein evaluations. This preference may be attributed to inconsistency between the training and inference stages: during training, the model is tasked with recovering full-length original structures, whereas in inference, it generates new structures and focuses on recovering motif regions only.

**Figure 4.**
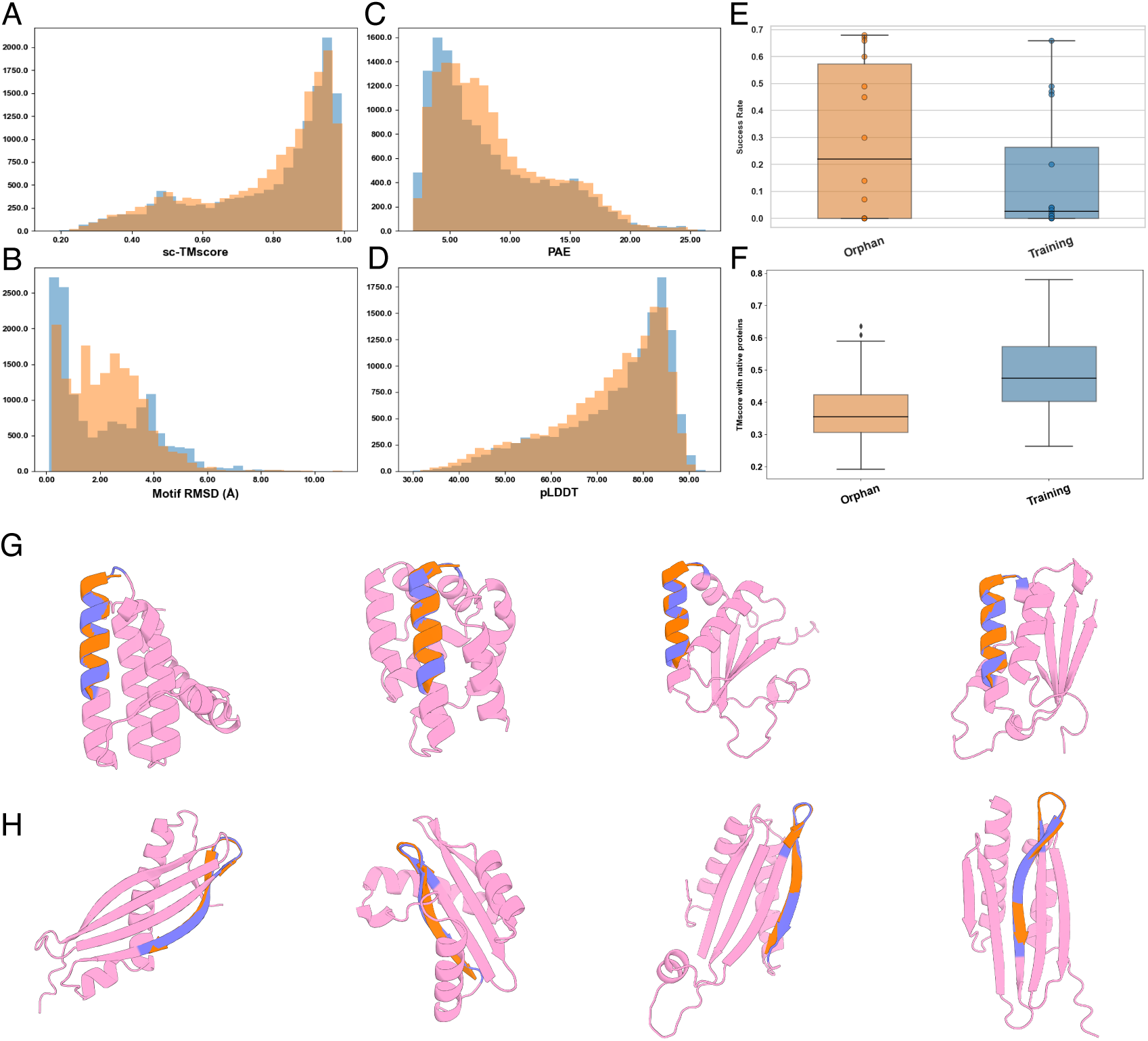
The robust performance of GPDL in orphan and training proteins. (**A-D**) sc-TMscore, motif RMSD, PAE and pLDDT distribution for orphan proteins (yellow) and proteins in the training set (blue). (**E**) Box plot and the scatter plot of the success rates for each orphan protein and the training protein. (**F**) Box plot of the TMscore for each successfully designed structure with respect to native structures. (**G-H)** Designed successful orphan protein (PDB ID: 7S5L) and training protein (PDB ID: 3FKA). Colors: native functional motif aligned to designed structure,orange; designed scaffold,pink; designed motif, purple.

Additionally, our generated orphan proteins possessed significantly lower TMscore with native structures compared to design backbone structures (**Fig. 4F**), indicating a high level of diversity among orphan proteins. We illustrated this with two examples: an orphan protein containing a helical motif (PDB ID: 7S5L, **Fig. 4G**) and a protein from the training set (PDB ID: 3FKA, **Fig. 4H**). Overall, GPDL is capable of generating a wide variety of protein backbones, including all-alpha structures, all-beta structures, and alpha-beta hybrid structures (**Fig. 4G-H and Fig. S13-S17**).

## Discussion

While our methods enhance computational efficiency and sample diversity, certain limitations persist. The capability for generating scaffolds around motifs consisting of a few number of residues has yet to be explored. For instance, it could be observed that most methods struggled to scaffold very short motifs ^28^, such as the case catalytic triad (PDB ID: 1QJG), which has only three discontiguous residues served as motif (**Fig. S4**). This is a representative problem for scaffolding active sites of enzymes, which plays a crucial role in protein engineering.

Moreover, the ability of controllable generation is one potential limitation of our present study. Improving control over the designed structures by integrating prior knowledge, such as secondary structures and small molecule positions, is one promising future direction of GPDL. Groundbreaking methods like AlphaFold3 ^26^ and RFdiffusion-AA^29^ have significantly advanced the modeling of complex macromolecule systems, incorporating various conditional inputs would benefit the current generation process of GPDL. In our optimization model, we experimented including the Van der Waals radius of ligands in the loss function to maintain the locations of binding molecules. However, while this penalty increased the distance between side-chain atoms and ligands, it did not significantly enhance generation accuracy (**Fig. S18**). Future work will focus on explicitly integrating functional information into the design process.

In addition, GPDL and most of the current methods only count for sequence and structure. However, the incorporation of sequences, structures and functions in the same space would benefit a lot for further applications. Notably, numerous high quality predicted protein structures from AlphaFold and ESMFold have made it possible to align structure embedding with current sequence-only language models. Furthermore, ESM3^11^ has emerged as a powerful tool for multimodal protein language modeling. Its prompt generation capabilities demonstrate that we can leverage more biological information to enhance guidance in protein design.

Besides, while GPDL reveals the promise of deep learning and fine-tuning in protein design, we solely validate our results *in silico*. Due to computational resource constraints, we did not extensively investigate the criteria for generation success. However, several experiments indicate that different metrics, such as TMscore or global RMSD as filters, indeed impact the success rates (**Fig. S19**). In this study, we follow the RFdiffusion benchmark protocol, setting the TMscore cutoff at 0.5 to evaluate additional properties and use ESMFold as the final filter. We observe that GPDL remains robust across different methods, including OmegaFold and the single-sequence version of AlphaFold2 (**Fig. S20**). This robustness is likely due to our use of ProteinMPNN for sequence redesign, aligning with findings in other research studies^30^. It is also worthwhile to note that further wet-lab validation of these computational methods remains essential as most of the artificial intelligence models try to please users by generating a computational reasonable protein but show deficient biophysical properties^31^.

Our exploration of motif scaffolding has sparked several research interests for future studies. At first, since one goal of backbone generation is to produce smaller, more stable proteins with specific functions by including other potential functional sites, it is essential to investigate the optimal sequence length. In this study, we investigated the quality of the backbones with different lengths. Our evaluation on the performance of backbones with varying sequence length on motif recapitulation and structure confidence represents an initial exploration of the problem of minimal backbone. Then, we observed a tendency for generated structures to reflect a bias toward the motif length. New methods are needed to precisely generate these short motif regions.

Our study underscores the significant impact of deep learning and protein language models on protein design. Many new powerful models derived from pre-trained language models have been a game changer in protein function annotation^32^, prediction of protein-protein interaction ^20^ and other biological applications^33^. Notably, numerous high quality predicted protein structures from AlphaFold ^34^ and ESMFold ^15^ have made it possible to add the current sequence-only language models ^11,35^.

In protein design, recent advances in deep learning tools have proven complementary to Rosetta ^36^, a widely used physics-based approach. Since many deep learning methods may lack robustness compared with physics-based approaches, several studies have successfully combined deep learning models with physical knowledge to achieve more plausible protein designs ^37,38^. This integrative approach leverages the strengths of both methodologies, resulting in improved design outcomes and a deeper understanding of protein engineering.

### Conclusion

In this study, we leveraged protein language models to develop a scalable motif-scaffolding method (GPDL) consisting of two lightweight and complementary modules. Our results demonstrate that GPDL can generate accurate and diverse scaffolds across various cases. Although recent deep generative models like diffusion models and flow matching shine in the protein design field, our methods and other works ^10,39^ proved that by converting structure prediction methods can still generate diverse and accurate proteins without heavily training resources.

## Materials and Methods

### Structure Seeding Module Model architectures

The model input consists of two parts: firstly, the language model is fed with active site sequences and “alanine padding mask” in ather positins, and the structure module takes the input of the distance matrix of active site residues in three-dimensional space. The distance matrix information includes the distances between each pair of amino acids for the 4 backbone atoms (N, CA, C, O) and 1 virtual atom, Virtual CB, totaling 25 distance measurements. The definition of the CB atom is as follows:

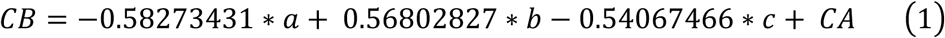

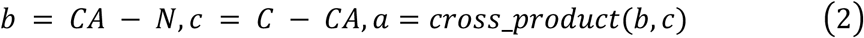

In this study, we adopt the Gaussian radial basis function to convert each distance into a 16-dimensional vector, following the same approach as employed in ProteinMPNN^25^. The *L_2_* distance between two distinct atoms, denoted as *m* and *n*, located in separate residues *i* and *j* is defined as follows:

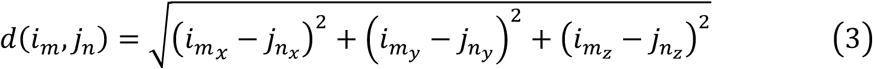

Then this distance will be transformed by 16 Gaussian radial functions (mean value from 2Å to 22Å)

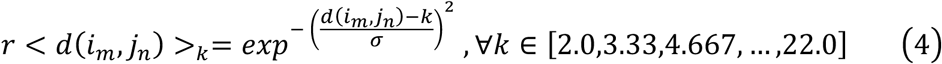

Main advantage of utilizing distance matrix is to achieve rotation and translation invariance and provides valuable three-dimensional protein prior knowledge. Any SE3 transformation *T* for the input structure *x* will not change the final output.

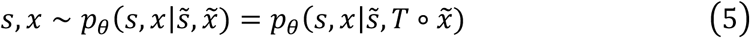

GPDL seeding module preserves most of the default configurations in ESMFold. The language model is esm2_t36_3B_UR50D version, folding trunk and structure module adopt esmfold_v1 version. We used an additive adapter to add the embedded distance matrix *d*_*ij*_ into the pair representation *z*_*ij*_ as follows. *W*, *b* represents the weight and bias in a single layer feed-forward network.

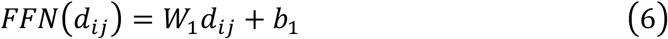

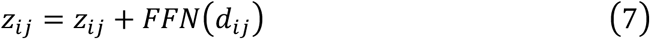

In the last recycle the single sequence representation *s_i_* will be projected into the probabilities of 20 amino acid. This module sequence predictor is defined as follow.

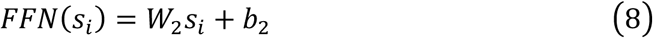

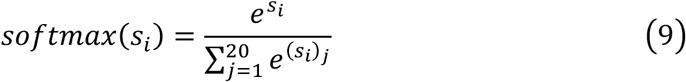

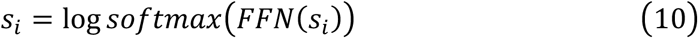

### Dataset and loss function

Seeding model employed the CATH4.3 ^40^ protein structure database as the dataset, ensuring that the redundancy among sequences did not exceed fourteen percent, which aligns with the ESM_IF ^21^ training dataset. During model training, for each protein sample, a random masking ratio between 0.3 and 0.6 was chosen, and accordingly, a portion of the sequences was masked with the alanine token, while some random chosen structural positions were masked with coordinates zero. As the sequence masking and structural masking are independent processes, we used a hybrid loss function to combine the global protein sequence cross-entropy loss and the frame-aligned point error (FAPE) for protein coordinates. The formulas are presented as follows:

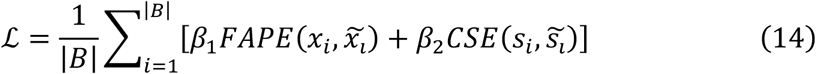

FAPE loss is initially proposed by AlphaFold2^1^ for protein structure prediction and demonstrate good performance in many downstream tasks such as protein diffusion models. The main idea is to align all the rigid groups and atoms between prediction results and target proteins. Here we only use one backbone rigid group and 3 atoms N, CA and C for each residue as follow. 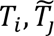 represents the backbone frame for the predicted residue *i* residue and the target residue *j*.

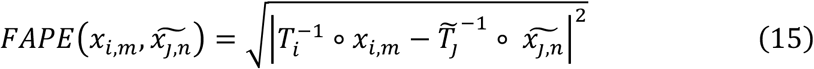

In this study, due to computational limitations we only fine-tune the structure module which is nearly 2 million parameters. We use Pytorch distributed data parallel and 4 graphics processing units (GPU) for the larger batch size. All the feature extraction from the PDB file is written by Biotite ^41,42^, Numpy ^43^ and Biopython ^44^.

### Structure Optimization Module Loss function

For a given functional motif, structure optimization module utilizes random flanking sequences as scaffolds and initiates the design by introducing the initial sequence into a cyclic optimization process. During the optimization process, the initial sequence undergoes the structure prediction module ESMFold. Since the prediction result at this stage stems from a random sequence, the initial prediction often yields an unfavorable disordered structure. To guide the design, a loss function is defined taking the desired property of the designs into account.

In this study, two key factors were considered when determining the loss function: the maintenance of motif conformation and the reliability of the generated backbone. The loss function is formulated as follows:

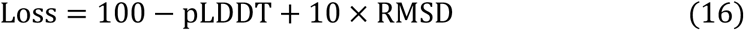

where the pLDDT (predicted Local Distance Difference Test) represents a measure of the confidence of the structure prediction, assessing the reliability of the designed backbone, and the Root Mean Square Deviation (RMSD) measures the root square mean deviation between the functional sites of the designed backbone and of the native structure.

By incorporating both pLDDT and RMSD in the loss function, the optimization process can effectively give protein backbones that not only retain the desired motif conformation but also exhibit higher reliability. This comprehensive approach ensures that the designed proteins are more likely to maintain their intended functionality while possessing a different structure to the native functional protein.

### MCMC Optimization

The optimization procedure guided by the loss function is based on MCMC (Markov Chain Monte Carlo) simulated annealing. After the initial round of structure prediction, random mutations are introduced into the scaffold sequence, and the structure prediction is performed on the updated sequences. The acceptance of updated sequence is according to the metropolis criterion:

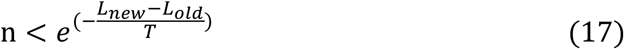

Where n is a value randomly sampled between 0 and 1, *L_old_* and *L_new_* are the losses calculated for the predicted results before and after the current round of mutation, and T is determined as:

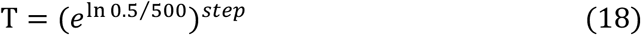

Where the step represents the current optimization round and the total number of optimization cycles for experiments in this study is determined as 1500. MCMC allows non-differentiable constraints to be a component of the loss function, making it applicated to other complex protein design problems beyond functional motif scaffolding.

As the loss function decreased, the designed backbone had optimized. After the final round of update and structure prediction, the optimized result was output. Through this interative optimization process, our model provided diverse backbones with high accuracy of the scaffolded functional motif.

### Sequence design and evaluation

While both methods were effective in generating the backbones and providing corresponding sequences, ProteinMPNN^12^ was used to refine the sequences corresponding to the acquired backbones, aiming to enhance the overall success rates during the subsequent evaluation process. During fixed backbone sequence design, all amino acid types within motif regions were fixed, identical to the original motifs.

For each case within the benchmark set, we used GPDL to generate 100 initial backbones. ProteinMPNN was then used to produce 10 sequences for each backbone. Afterwards, we used ESMFold to predict the 3D structures of each sequence and calculated the structural similarity with initially generated backbones, of which the procedure is commonly referred as designable evaluation or self-consistency test. We consider three metrics as designability definition: Motif reconstruction, with RMSD between ESMFold predicted motif and native motif less than 1.0 Å; backbone reconstruction, with TMscore between ESMFold predicted structure and initially generated backbone larger than 0.5; and a structure prediction confidence metric, where the predicted aligned error (PAE) value less than 5 Å. Under this criteria, a backbone is considered designable if at least one out of 10 ProteinMPNN generated sequences meet all of these criteria. Notably, our definition slightly deviates from the original method^5^, which employs a single sequence AlphaFold prediction for self-consistency. We hypothesized that the current state-of-the-art single sequence prediction model ESMFold shows better performance than AlphaFold2 with no MSA information. By collectively applying these evaluation metrics, we gain valuable insights into the quality and reliability of the generated protein structures, enabling a comprehensive assessment of their performance.

### Design on metal-binding proteins

The sequences of metal-binding proteins 1BCF and 1PRW were designed using ProteinMPNN and then predicted using the online server of AlphaFold3. All-atom MD simulations were carried out using the AMBER 22 package ^45^. The Amber *ff03CMAP* force field ^46^ and TIP4PEW water model ^47^ were used, with parameters of metal ions generated using *gaff2* force field. All systems were neutralized and solvated in water with cubic boxes ensuring a minimum distance of 10 Å between the box edges and the protein. All bonds involving hydrogen atoms were constrained with the SHAKE algorithm. The particle mesh Ewald (PME) algorithm was used to calculate long-range electrostatic interactions. Initial structures were relaxed with 10,000 steps of minimization, then subjected to heating for 50 ps from 0 K to 300 K and equilibration for 100 ps in NPT ensemble with PMEMD. The simulation was later conducted with the temperature set to 300K.

## ACKNOWLEDGMENTS

This work was supported by Shanghai Municipal Science and Technology Major Project, partially by SJTU Kunpeng&Ascend Center of Excellence, the Center for HPC at Shanghai Jiao Tong University, and the National Key Research and Development Program of China (2023YFF1205102 and 2020YFA0907700), the Fundamental Research Funds for the Central Universities (YG2023LC03), and the National Natural Science Foundation of China (32171242).

## Code Availability

The source code of GPDL is available at https://github.com/sirius777coder/GPDL.

