## Supplementary Information for "Protein Language Model Supervised Scalable Approach for Diverse and Designable Protein Motif-Scaffolding with GPDL"

Bo Zhang#, Kexin Liu#, Zhuoqi Zheng#, Junjie Zhu, Zhengxin Li, Yunfeiyang Liu, Junxi Mu, Ting Wei*, Hai-Feng Chen*
State Key Laboratory of Microbial metabolism, Joint International Research Laboratory of Metabolic & Developmental Sciences, Department of Bioinformatics and Biostatistics, National Experimental Teaching Center for Life Sciences and Biotechnology, School of Life Sciences and Biotechnology, Shanghai Jiao Tong University, Shanghai, 200240, China

***Corresponding Author**
**Hai-Feng Chen (Full Professor)**
State Key Laboratory of Microbial metabolism, Joint International Research Laboratory of Metabolic & Developmental Sciences, Department of Bioinformatics and Biostatistics, National Experimental Teaching Center for Life Sciences and Biotechnology, School of Life Sciences and Biotechnology, Shanghai Jiao Tong University, Shanghai, 200240, China

**Notes**

The authors declare that there is no conflict of interest.

**Author Contributions**

#These authors contributed equally to this work.

**Table S1. Model performance in all design sequences.** Diversity represents the average number of unique designable clusters across 24 cases. The overall success rate is calculated as the ratio of all successful designs to the total number of designs (24,000). The success rate is further averaged per case to account for the fraction of successful backbones average in each target.

| Method | Unique Designable Cluster | Success Rate (Sequence) | Success Rate (Backbone) |
|---|---|---|---|
| **GPDL-1500** | **7.677** | 0.242 | 0.371 |
| **GPDL-800** | 6.468 | 0.196 | 0.302 |
| **GPDL (No prompt)** | 6.417 | 0.266 | 0.399 |
| **RFdiffusion** | 5.750 | **0.267** | **0.445** |
| **RFdesign-Inpainting** | 0.708 | 0.194 | 0.311 |
| **RFdesign-Hallucination** | 2.250 | 0.128 | 0.223 |
| **TDS** | 5.583 | 0.093 | 0.257 |
| **Chroma** | 1.833 | 0.075 | 0.161 |

**Table S2. Model success rate in 24 cases.** * indicates that the highest success rate among GPDL-1500, GPDL-800, and GPDL-No Prompt was selected for GPDL. † denotes that all successful 5YUI designs were generated by GPDL-5000. ‡ denotes that the successful 1QJG design was generated by GPDL (no prompt).

| | GPDL* | RFdiffusion | RFdesign-Inpainting | RFdesign-Hallucination | Chroma | TDS |
|---|---|---|---|---|---|---|
| **1BCF** | 0.17 | **1** | 1 | 0.37 | 0.42 | 0.01 |
| **1PRW** | **1** | 0.99 | 1 | 0.25 | 0.84 | 0.44 |
| **1QJG** | 0.01‡ | 0 | 0 | 0.24 | 0 | 0.07 |
| **1YCR** | 0.88 | **0.99** | 0.77 | 0.98 | 0.55 | 0.6 |
| **2KL8** | 0.8 | 0.59 | 0.8 | 0.07 | 0.92 | 0.21 |
| **3IXT** | **1** | 0.91 | 1 | 0.92 | 0.66 | 0.53 |
| **4JHW** | 0 | 0 | 0 | 0 | 0 | 0 |
| **4ZYP** | **0.85** | 0.67 | 0.08 | 0.44 | 0.24 | 0.02 |
| **5IUS** | 0.04† | 0.01 | 0.2 | 0 | 0 | 0 |
| **5TPN** | 0.32 | **0.76** | 0.43 | 0.04 | 0 | 0 |
| **5TRV_long** | 0.09 | **0.54** | 0 | 0.04 | 0 | 0.02 |
| **5TRV_medium** | 0.07 | 0.35 | 0 | 0.02 | 0 | 0.42 |
| **5TRV_short** | 0.04 | 0.2 | 0 | 0 | 0 | 0.22 |
| **5WN9** | **0.13** | 0 | 0 | 0 | 0.02 | 0 |
| **5YUI** | 0 | 0 | 0 | 0 | 0 | 0 |
| **6E6R_long** | 0.6 | **0.78** | 0 | 0.64 | 0.03 | 0.53 |
| **6E6R_medium** | **0.69** | 0.52 | 0 | 0.41 | 0.01 | 0.64 |
| **6E6R_short** | **0.73** | 0.19 | 0.1 | 0.62 | 0.06 | 0.51 |
| **6EXZ_long** | 0.33 | **0.9** | 0.18 | 0.22 | 0.03 | 0.19 |
| **6EXZ_medium** | 0.3 | 0.74 | 0.81 | 0.05 | 0.04 | 0.86 |
| **6EXZ_short** | 0.27 | 0.07 | 0.83 | 0.02 | 0.04 | 0.85 |
| **7MRX_128** | **0.41** | 0.14 | 0 | 0.01 | 0 | 0 |
| **7MRX_60** | **0.66** | 0.14 | 0.25 | 0 | 0 | 0.02 |
| **7MRX_85** | **0.58** | 0.18 | 0.01 | 0 | 0 | 0.03 |

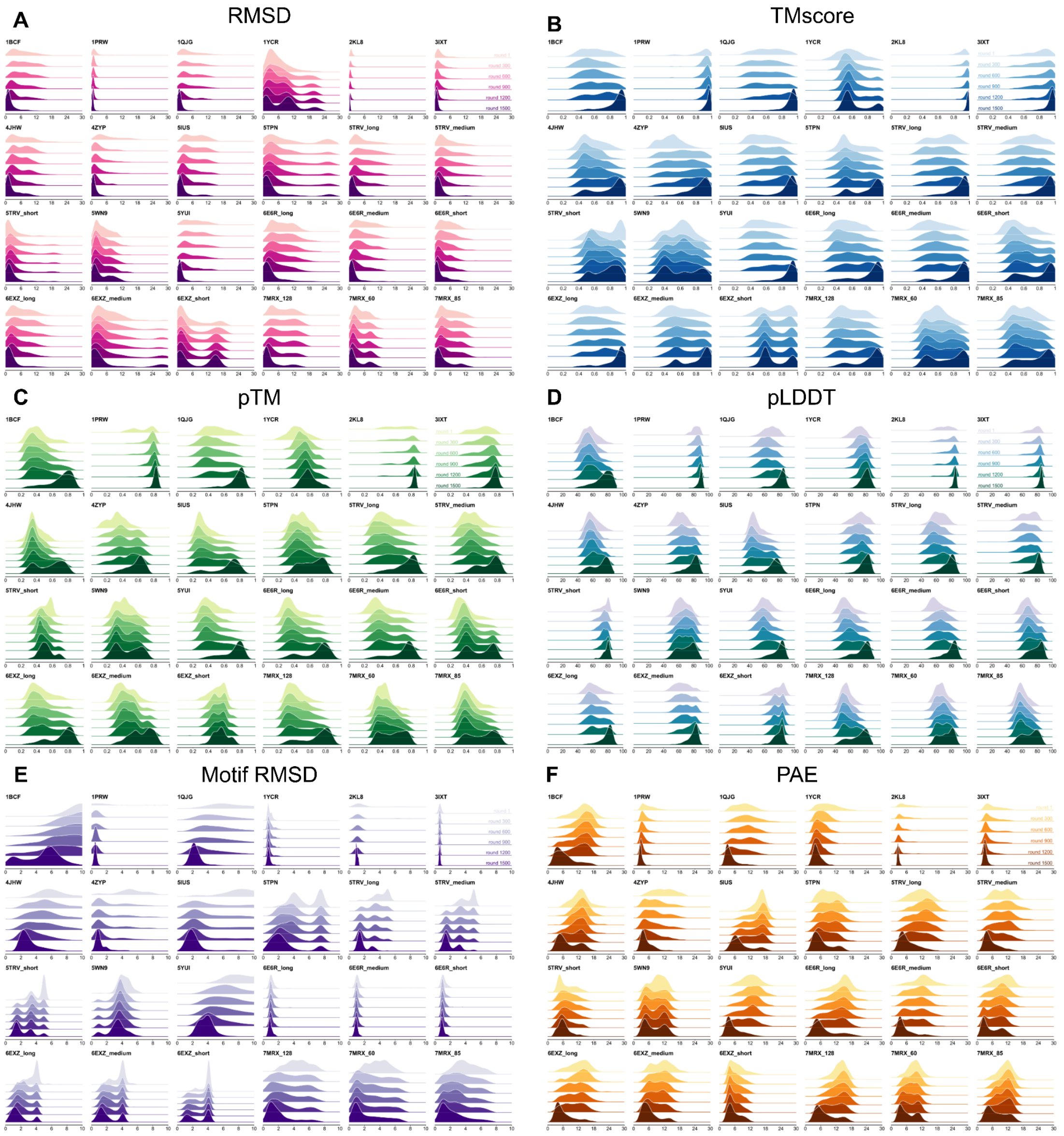


**Figure S1. Optimization trajectories for 24 cases. Metrics displayed here are calculated under the scenario of self-consistency test.** (**A**) RMSD over different cases and time steps. (**B**) scTM-score over different cases and time steps. (**C**) pTM score over different cases and time steps. (**D**) pLDDT score over different cases and time steps. (**E**) Motif RMSD over different cases and time steps. (**F**) Predicted Alignment Error (PAE) over different cases and time steps.

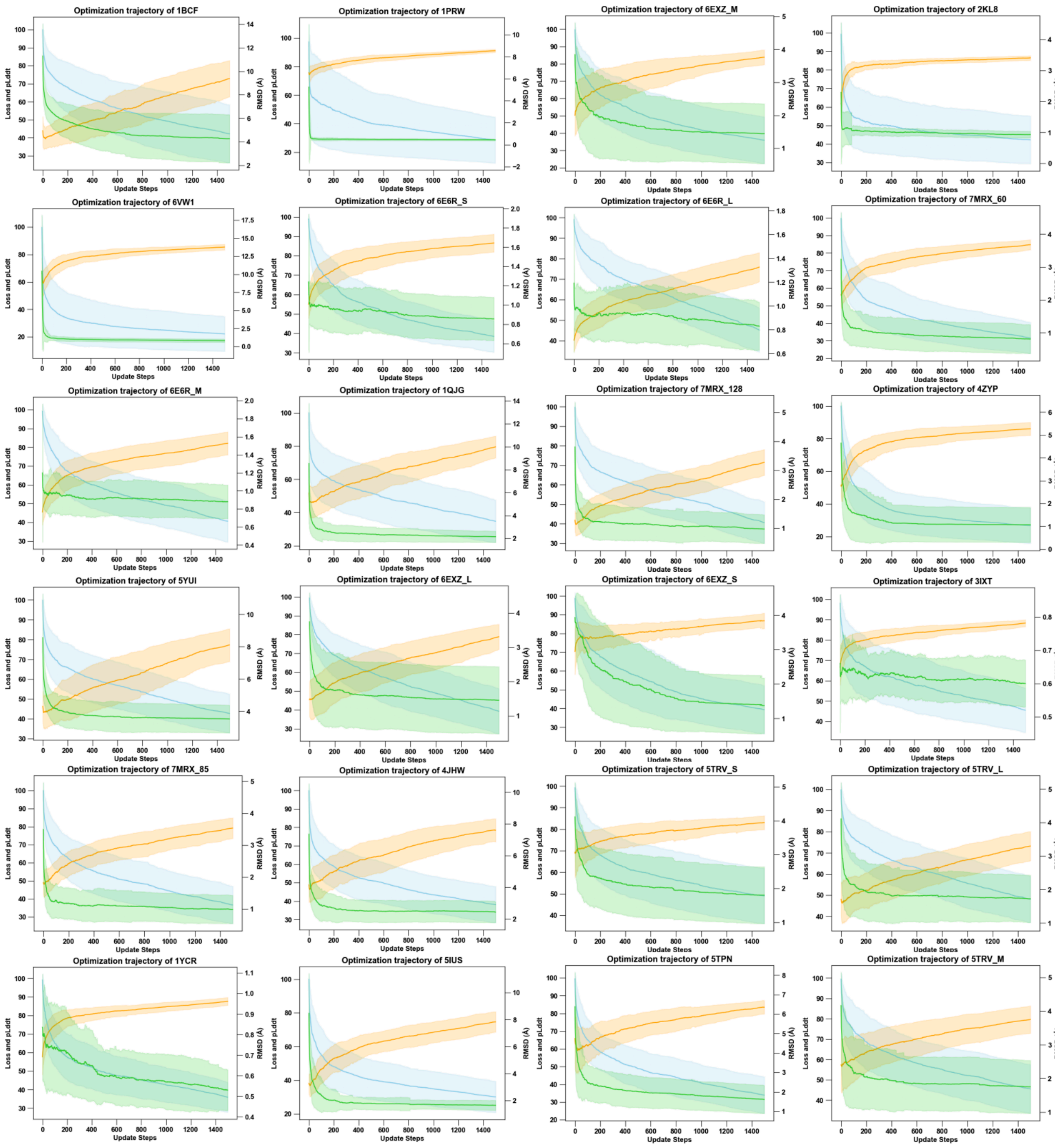


**Figure S2. Loss curve of 24 cases during optimization.**

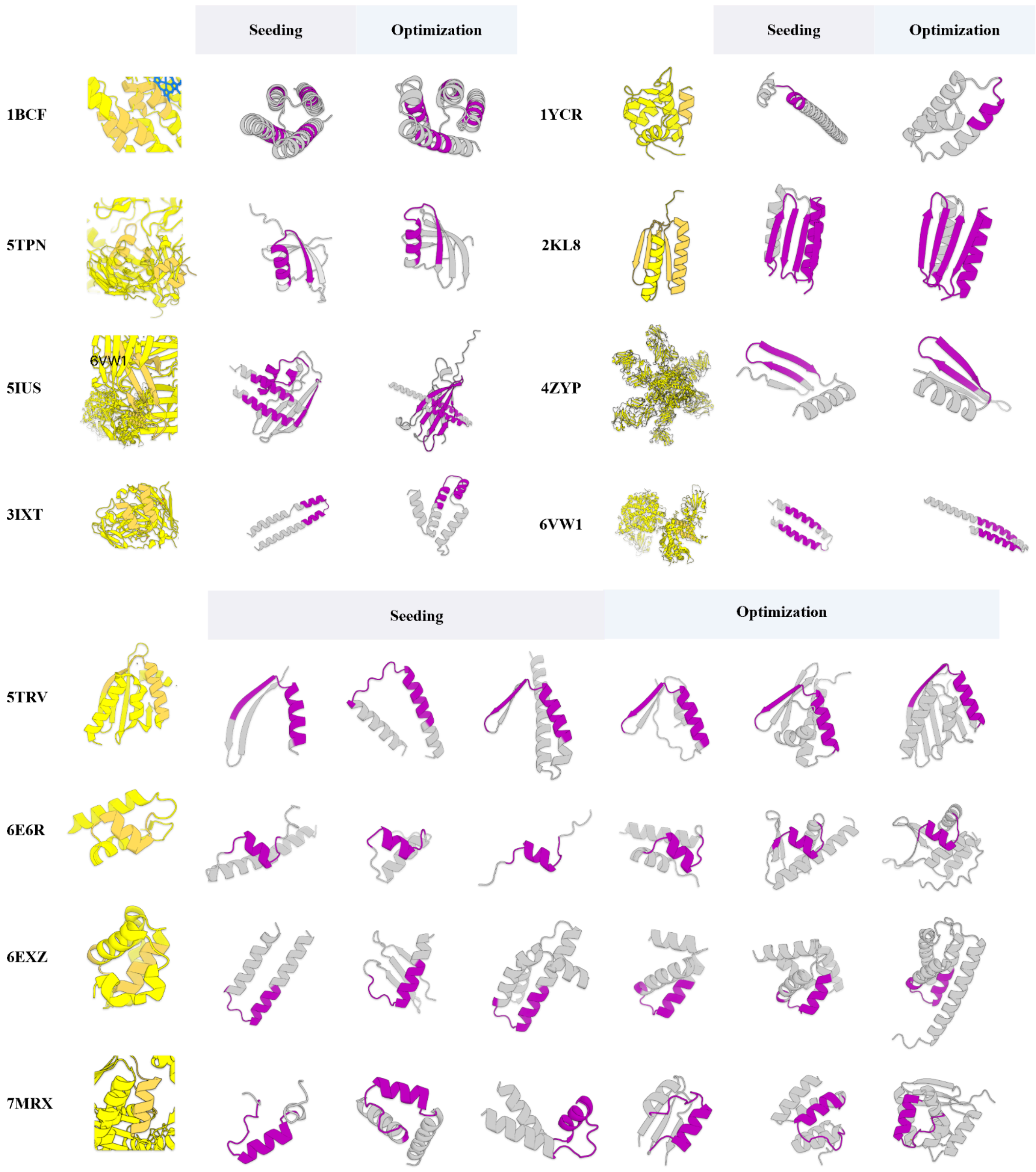


**Figure S3. designs by seeding module or optimization module.**

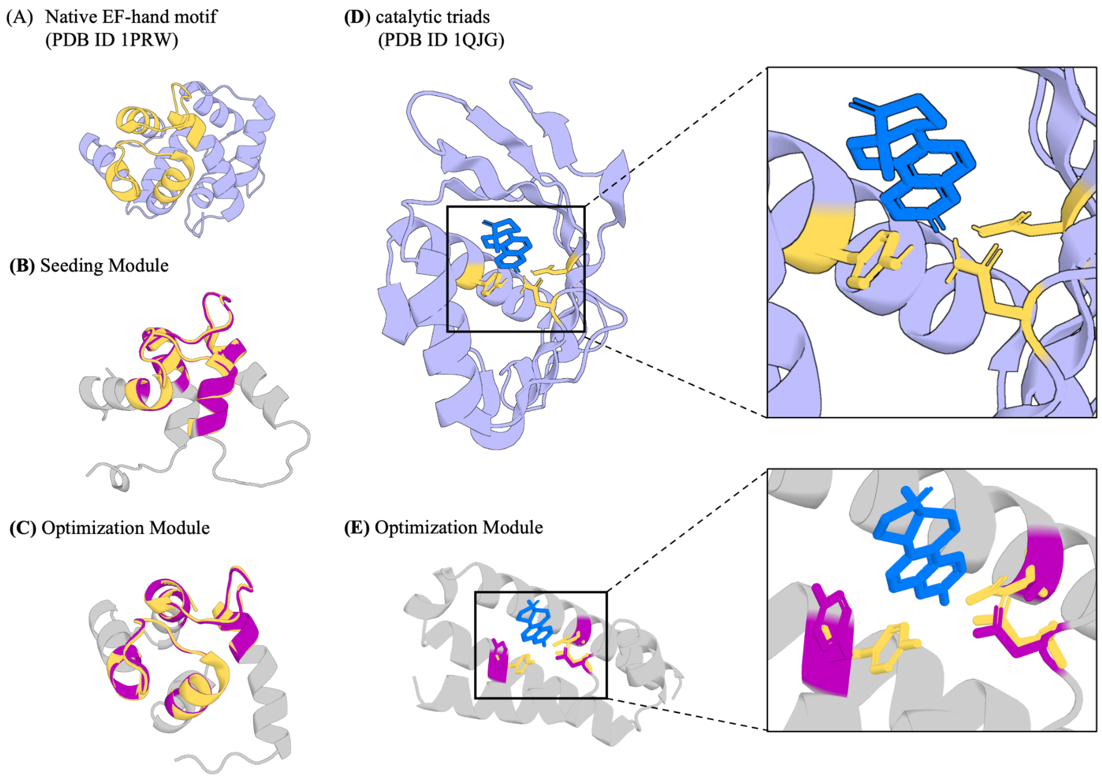


**Figure S4**. **Design of metal-coordinating protein and enzymes.** (A-C) 1PRW metal binding functional sites. (A) Native protein PDB ID 1PRW and the EF-hand motif chain A residues 16 to 35 and 52 to 71. (B) Seeding EF-hand motif. Accuracy : motif RMSD = 0.398Å (with native motif), prediction confidence: PAE = 4.42Å, self-consistency RMSD = 1.78Å. (C) Optimization EF-hand motif. Accuracy : motif RMSD = 0.33Å (with native motif), prediction confidence: PAE = 2.76Å. self-consistency RMSD = 0.62Å. Colors: native protein scaffold, yellow; native functional motif, yellow-orange; seeding scaffold, gray; seeding motif, purple. (D and E) 1QJG catalytic triads scaffolding.(D) Native protein PDB ID 1QJG and the motif A residues 16 to 35 and 52 to 71. (E) Optimization catalytic triads. Accuracy : motif RMSD = 0.93Å (with native motif), prediction confidence: PAE = 2.66Å, self-consistency RMSD = 0.62Å.Colors: native protein scaffold, yellow; native functional motif, yellow-orange; seeding scaffold, gray; seeding motif, purple.

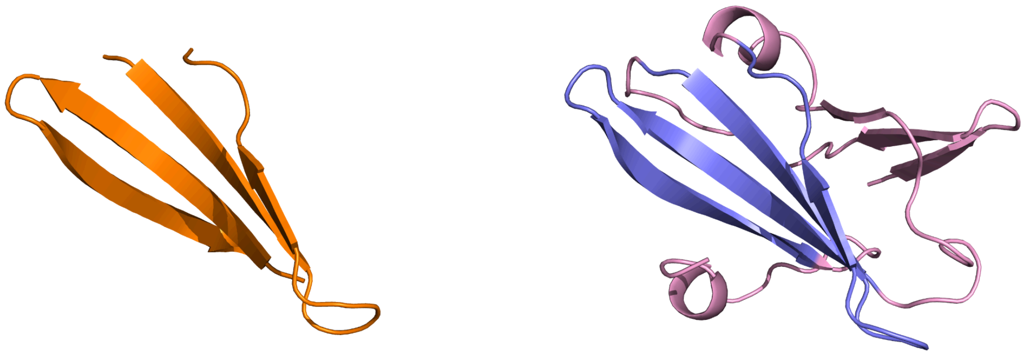

**Figure S5. GPDL generated successful scaffolds for 5IUS**. The native motif targeting PD-L1 (PDB id: 5IUS) is shown in orange, design motifs in blue, and design scaffolds in pink. This design is generated by GPDL after 5000 round optimization with prompt.

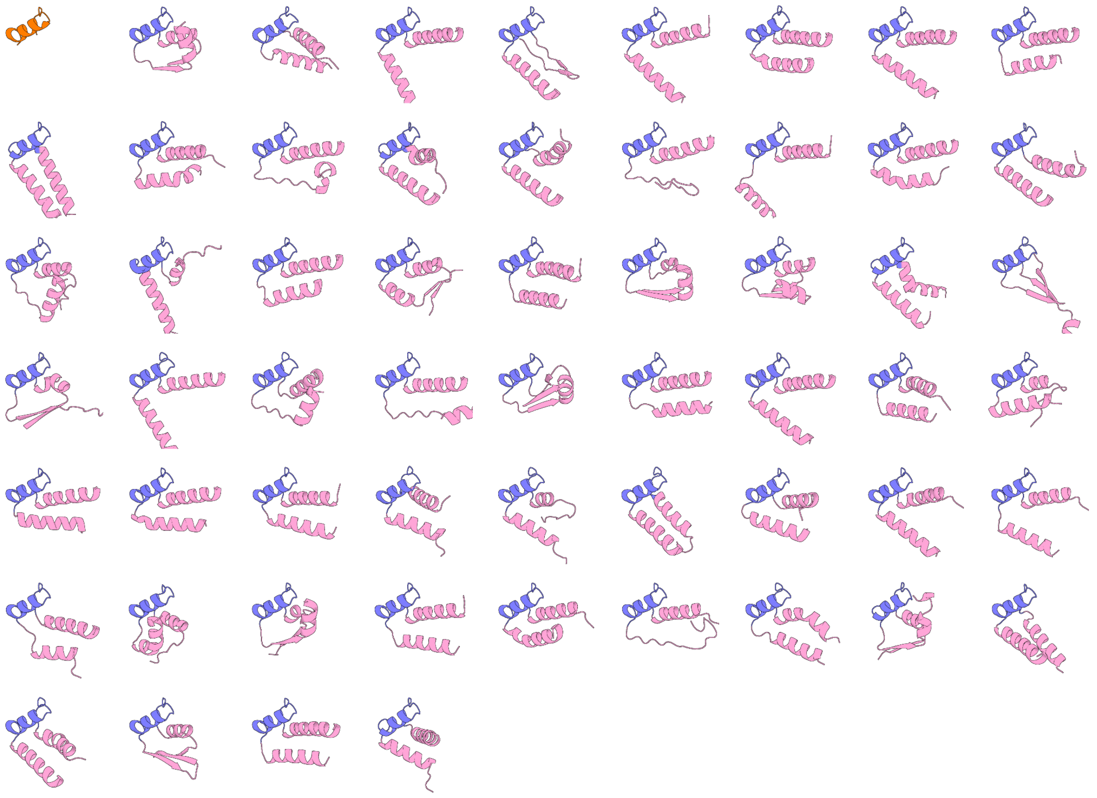

**Figure S6. GPDL generated scaffolds for 7MRX_60**. The native motif from PDB ID 7MRX is shown in orange, design motifs in blue, and design scaffolds in pink.

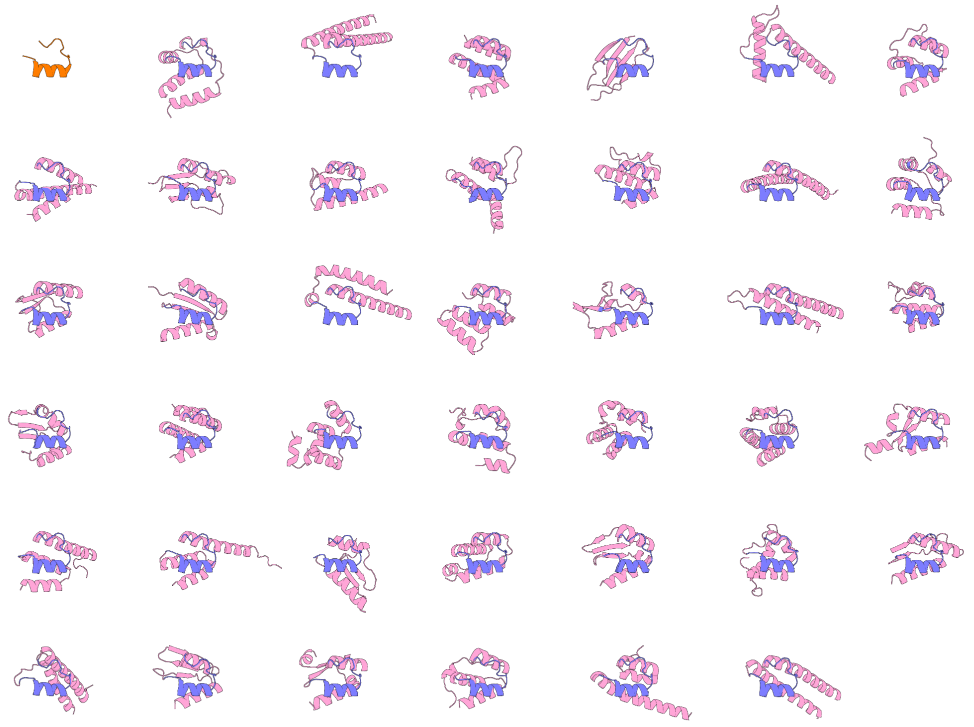

**Figure S7. GPDL generated scaffolds for 7MRX_85**. The native motif from PDB ID 7MRX is shown in orange, design motifs in blue, and design scaffolds in pink.

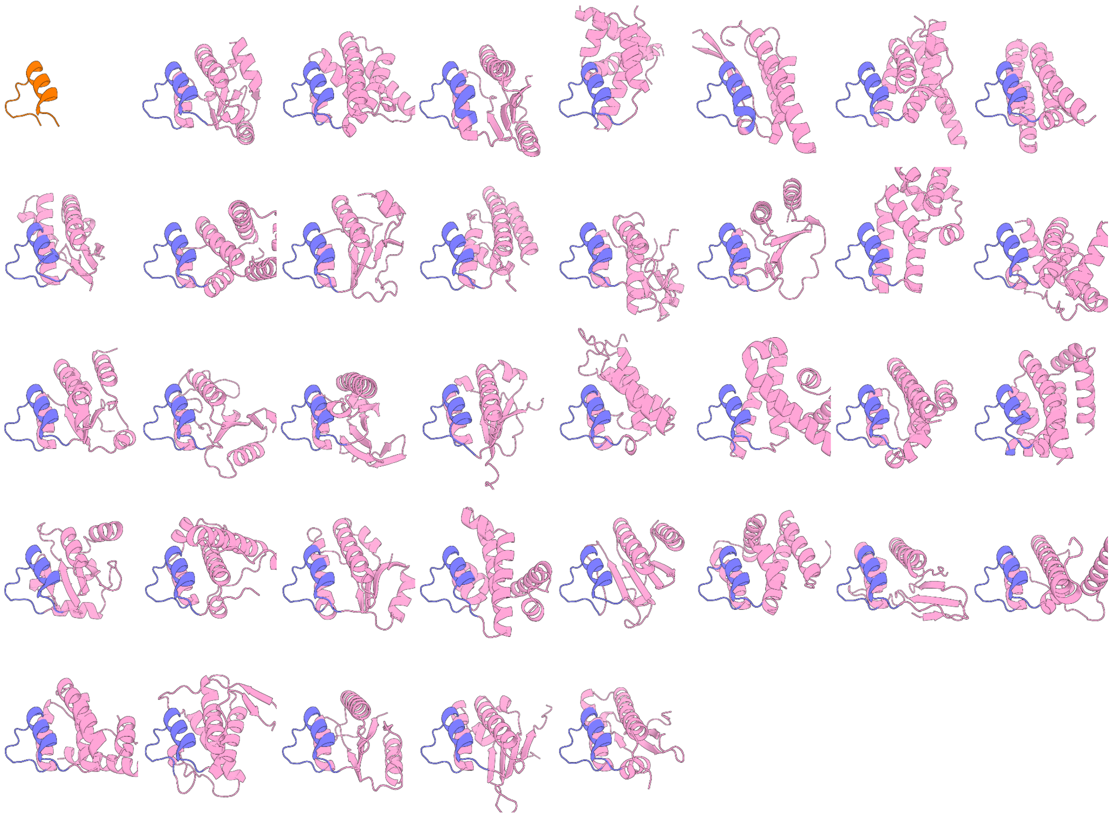

**Figure S8. GPDL generated scaffolds for 7MRX_128**. The native motif from PDB ID 7MRX is shown in orange, design motifs in blue, and design scaffolds in pink.

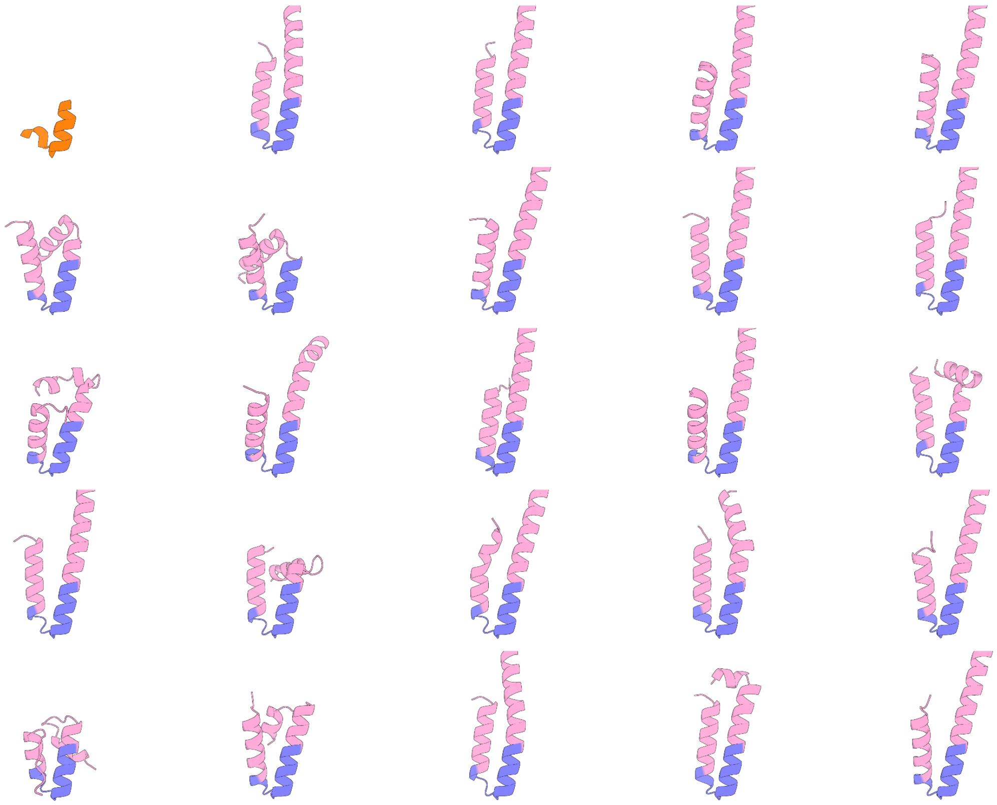

**Figure S9. GPDL generated scaffolds for 6EXZ_short**. The native motif from PDB ID 6EXZ is shown in orange, design motifs in blue, and design scaffolds in pink.

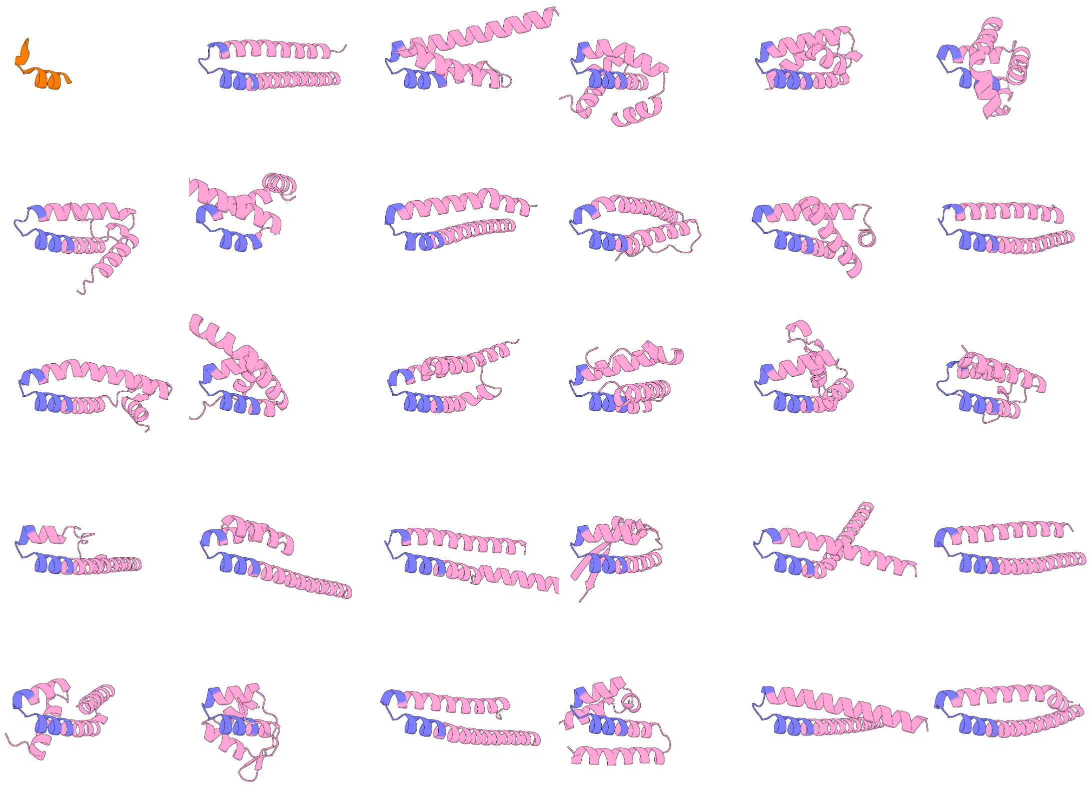

**Figure S10. GPDL generated scaffolds for 6EXZ_medium**. The native motif from PDB ID 6EXZ is shown in orange, design motifs in blue, and design scaffolds in pink.

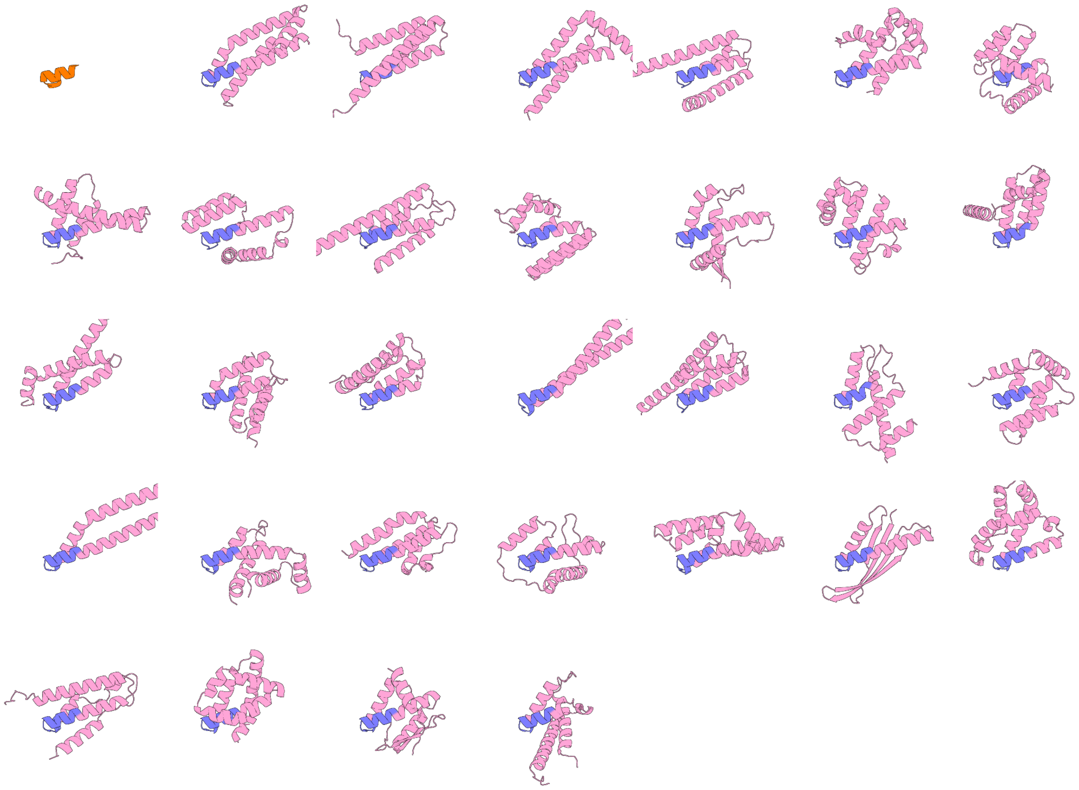

**Figure S11. GPDL generated scaffolds for 6EXZ_long.** The native motif from PDB ID 6EXZ is shown in orange, design motifs in blue, and design scaffolds in pink.

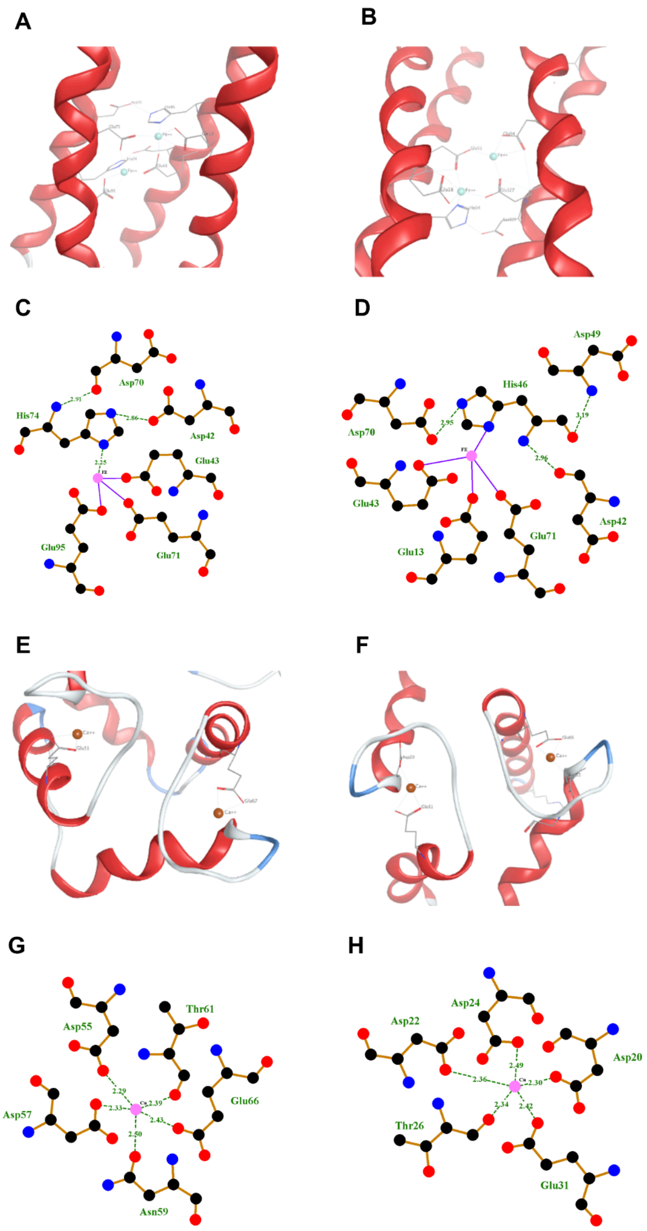


**Figure S12. Metal binding motif analysis.** (A, B) MOE plots of the native metal-binding motif in PDB ID 1BCF and the corresponding designed protein scaffold containing the 1BCF motif. (C, D) Ligplot diagrams of the designed protein scaffold with the 1BCF motif, showing two iron ions. (E, F) MOE plots of the native metal-binding motif in PDB ID 1PRW and the designed protein scaffold containing the 1PRW motif. (G, H) Ligplot diagrams of the designed protein scaffold with the 1PRW motif, showing two calcium ions. All designed structures were predicted by AlphaFold3.

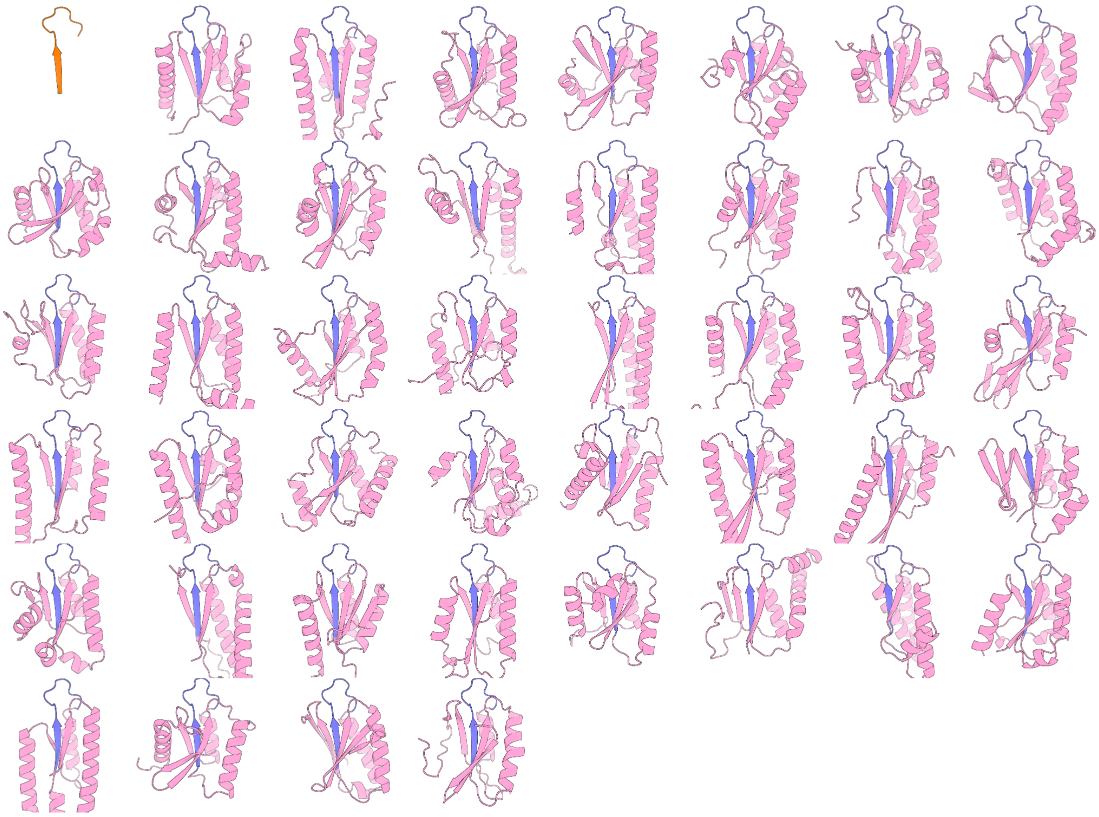

**Figure S13. GPDL generated scaffolds for the training set protein 4M1T.**
The native beta-strand motif from PDB ID 4M1T is shown in orange, design motifs in blue, and design scaffolds in pink.

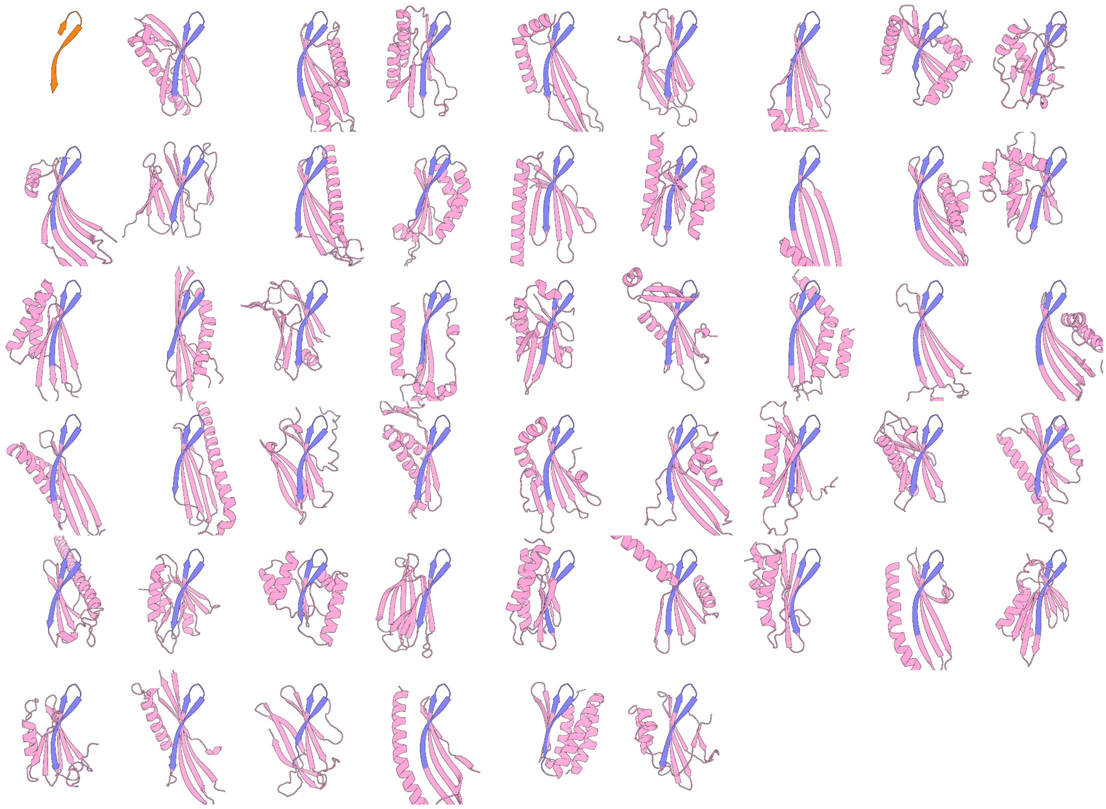

**Figure S14. GPDL generated scaffolds for the training set protein 3FKA.**
The native beta-strand motif from PDB ID 3FKA is shown in orange, design motifs in blue, and design scaffolds in pink.

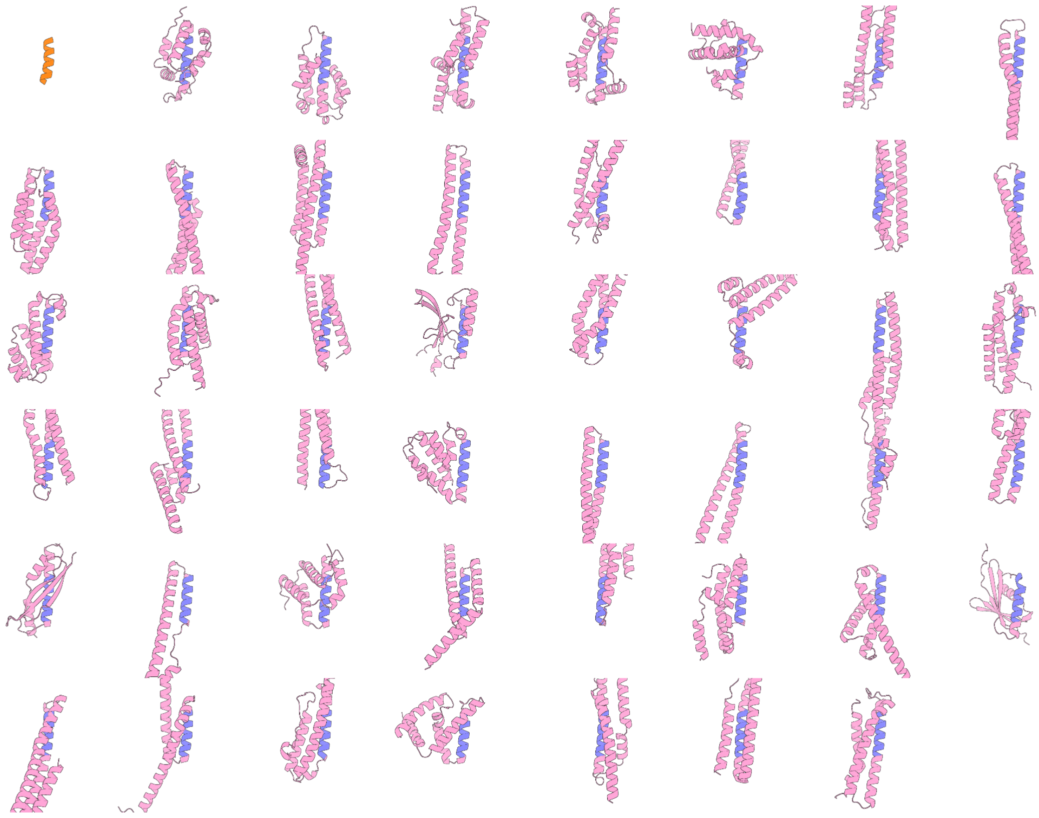

**Figure S15. GPDL generated scaffolds for the orphan set protein 7TJL.**
The native beta-strand motif from PDB ID 7TJL is shown in orange, design motifs in blue, and design scaffolds in pink.

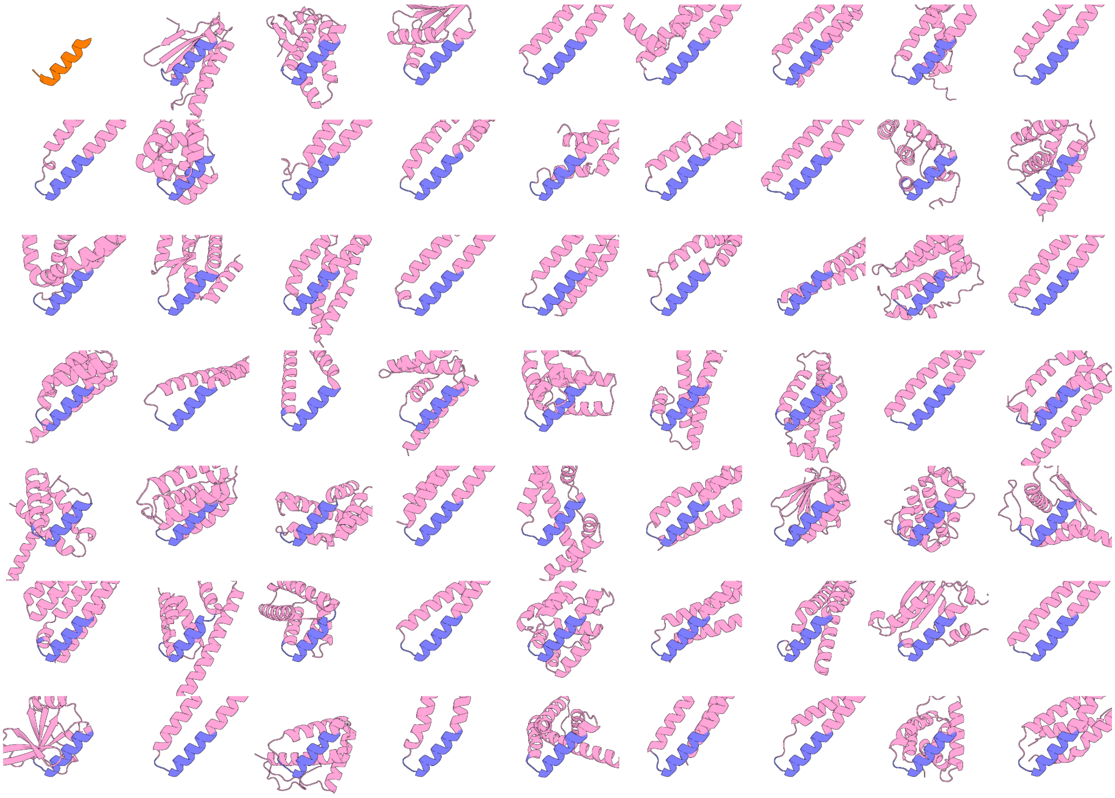

**Figure S16. GPDL generated scaffolds for the orphan set protein 7S5L.**
The native beta-strand motif from PDB ID 7TJL is shown in orange, design motifs in blue, and design scaffolds in pink.

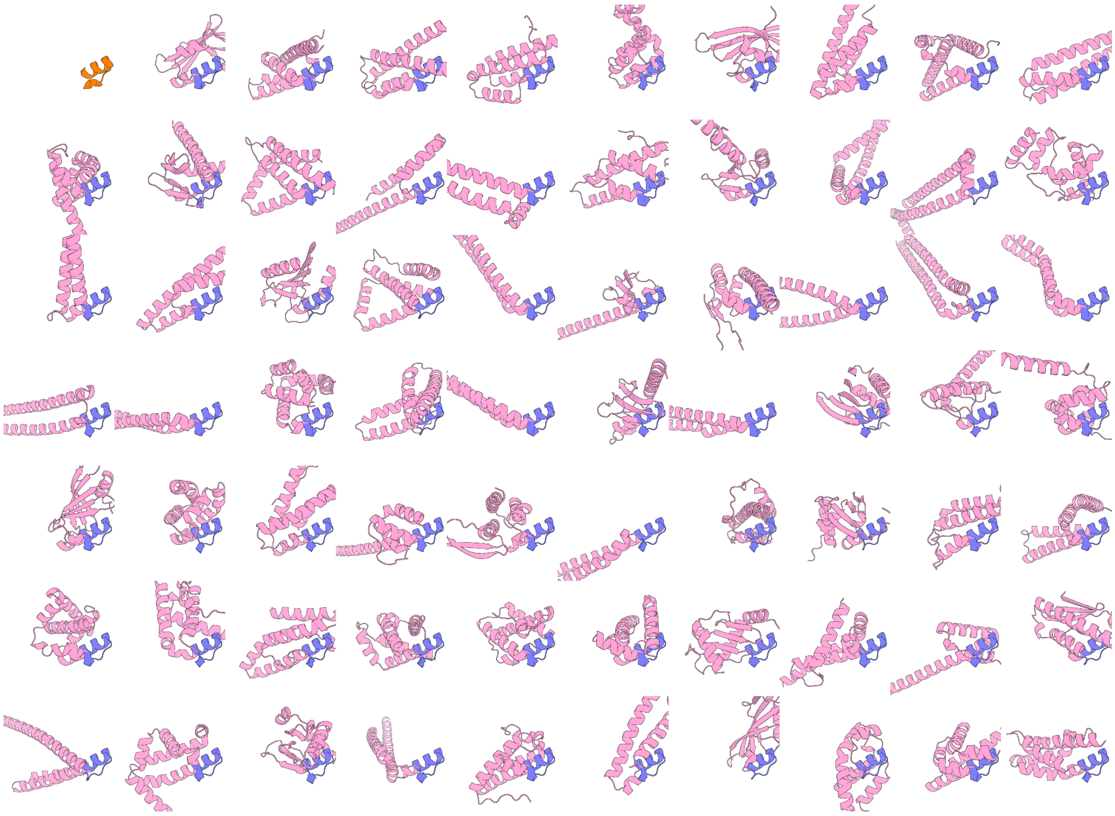

**Figure S17. GPDL generated scaffolds for the orphan set protein 7KUW.**
The native beta-strand motif from PDB ID 7KUW is shown in orange, design motifs in blue, and design scaffolds in pink.

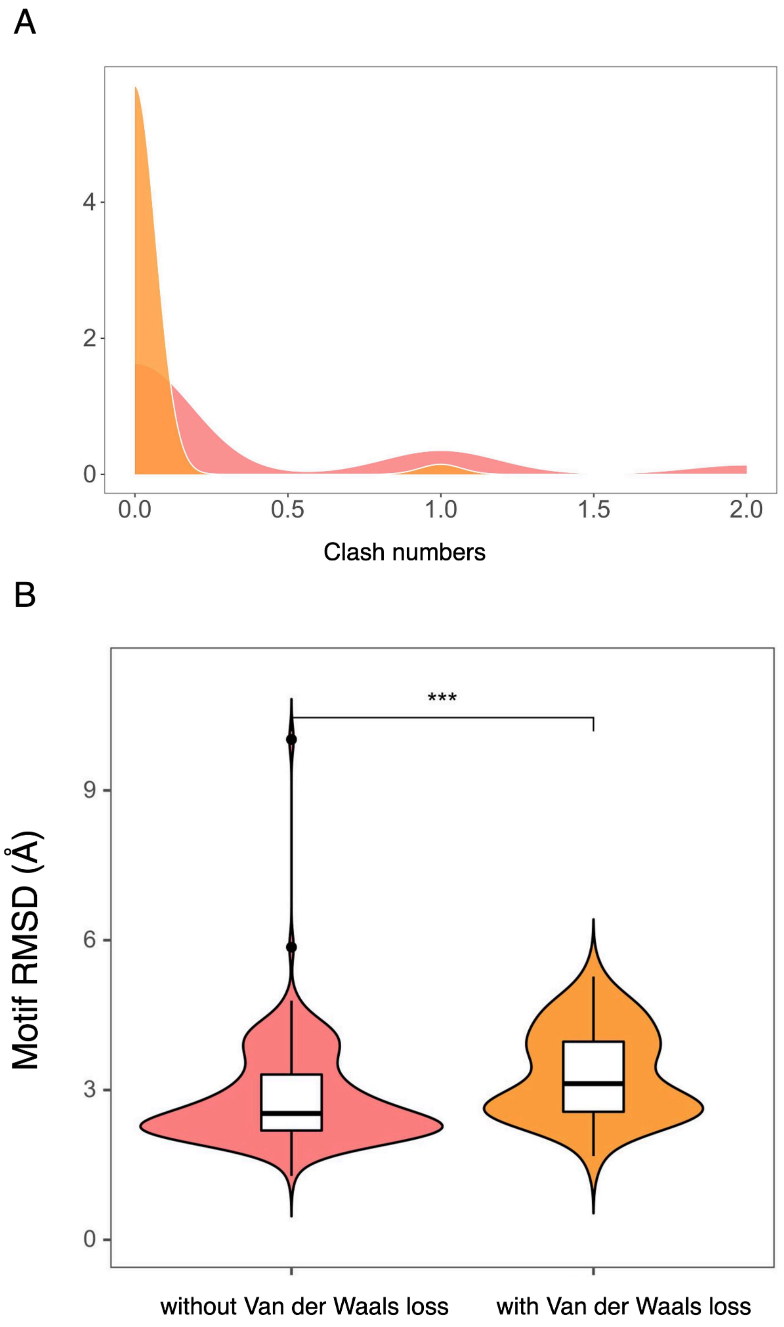


**Figure S18. Effect of incorporating Van der Waals loss during optimization.** (A) Comparison of atomic clashes with and without Van der Waals loss in the optimization module. (B) Motif RMSD with and without incorporation of Van der Waals loss in the optimization module.

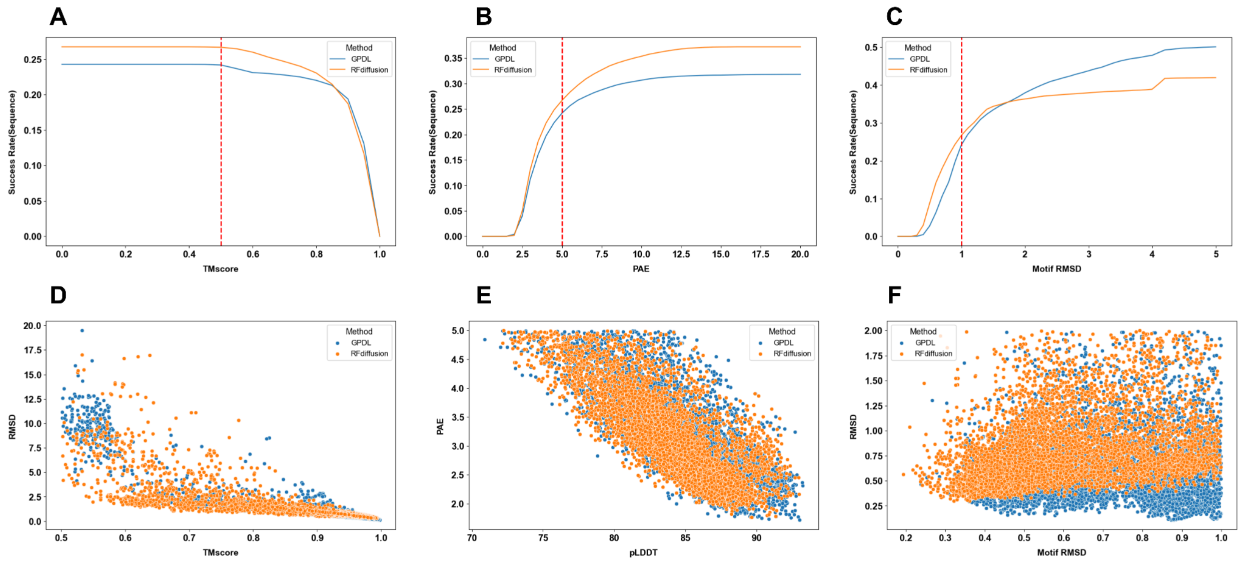


**Figure S19. GPDL and RFdiffusion success rate cutoff.** (A-C) Different metrics cutoff and the corresponding overall success rate, and let other metrics be fixed. (D-F) Pairwise metric plot for GPDL successful designs and RFdiffusion successful designs

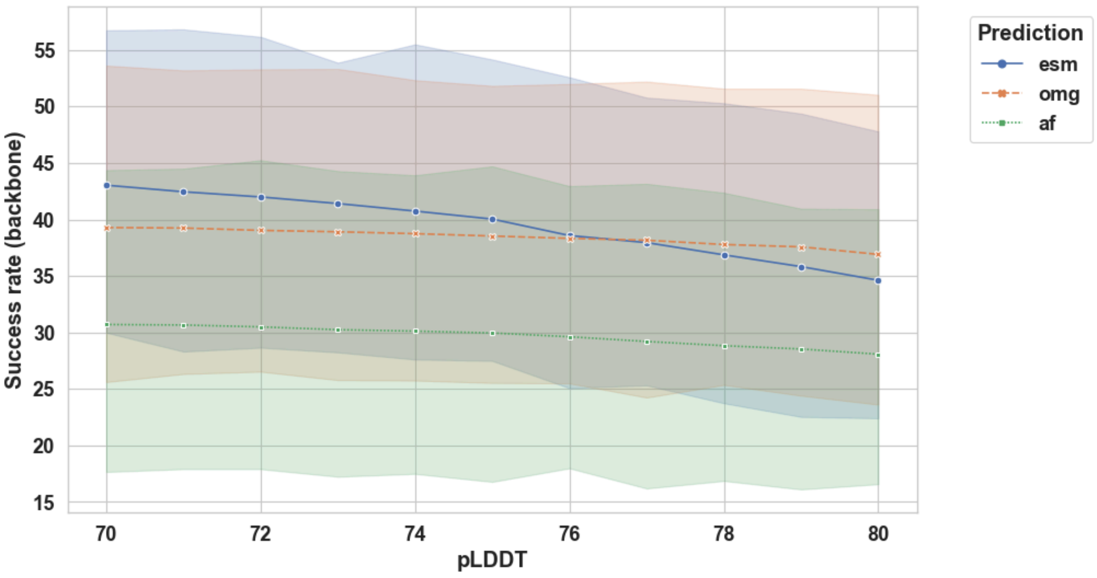


**Figure S20. Success rate of GPDL in different pLDDT cutoffs with different refolding methods.**
